# Crystal structure of full-length cytotoxic necrotizing factor CNF_Y_ reveals molecular building blocks for intoxication

**DOI:** 10.1101/2020.04.07.029181

**Authors:** Paweena Chaoprasid, Peer Lukat, Sabrina Mühlen, Thomas Heidler, Emerich-Mihai Gazdag, Shuangshuang Dong, Wenjie Bi, Christian Rüter, Marco Kirchenwitz, Anika Steffen, Lothar Jänsch, Theresia E. B. Stradal, Petra Dersch, Wulf Blankenfeldt

## Abstract

Cytotoxic necrotizing factors (CNFs) are bacterial single-chain exotoxins that modulate cytokinetic/oncogenic and inflammatory processes through activation of host cell Rho GTPases. To achieve this, they are secreted, bind surface receptors to induce endocytosis and translocate a catalytic unit into the cytosol to intoxicate host cells. A three-dimensional structure that provides insight into the underlying mechanisms is still lacking. Here, we determined the crystal structure of full-length *Yersinia pseudotuberculosis* CNF_Y_. CNF_Y_ consists of five domains (D1-D5), and by integrating structural and functional data we demonstrate that D1-3 act as export and translocation module for the catalytic unit (D4-5) or fused β-lactamase reporter proteins. We further found that domain D4, which possesses structural similarity to ADP-ribosyl transferases, but had no equivalent catalytic activity, changed its position to interact extensively with D5 in the crystal structure of the free D4-5 fragment. This liberates D5 from a semi-blocked conformation in full-length CNF_Y_, leading to higher deamidation activity. Finally, sequence comparisons identified the CNF translocation module in many uncharacterized bacterial proteins, suggesting its usability as a universal drug delivery tool.

## Introduction

Amongst the plethora of traits developed by pathogenic bacteria to establish infections, toxins play the most prominent role, since they are responsible for the majority of clinical symptoms (Popoff, 2005). Many bacterial exotoxins are key virulence factors that target different functions of host cells to break barriers, improve access to nutrients, defeat immune responses and promote bacterial dissemination within and among hosts.

The cytotoxic necrotizing factors (CNFs) belong to a class of bacterial exotoxins that deamidate a glutamine (Q61 or Q63) in the active site (switch II region) of host cell proteins belonging to the small Rho GTPase family, i.e. RhoA, Rac1 and Cdc42 (Flatau *et al*, 1997; Schmidt *et al*, 1997; Knust & Schmidt, 2010). This locks these key regulators in their active state, causing a multitude of downstream effects that are most readily observed as alterations of the actin cytoskeleton or perturbations of other cellular processes including phagocytosis, cell proliferation (multinucleation), reactive oxygen species production, and the release of anti-apoptotic and pro-inflammatory factors (Fabbri *et al*, 2013; Hodge & Ridley, 2016; Ho *et al*, 2018). The consequences of these effects are changes of innate immune responses and tissue damage, leading to the development of acute disease symptoms (Knust & Schmidt, 2010; Schweer *et al*, 2013; Diabate *et al*, 2015; Cavaillon, 2018; Heine *et al*, 2018).

CNFs are found in several pathogenic bacteria, predominantly in pathogenic *Escherichia coli*, but also in *Yersinia pseudotuberculosis*, *Shigella* species, *Salmonella enterica,* as well as in *Moritella viscosa* and *Photobacterium damselae*, pathogens of economically important fish (Fig EV1) (Morgan *et al*, 2019). In addition, CNFs show local homology to other toxins such as the dermonecrotizing toxin DNT of *Bordetella pertussis* and the PMT toxin of *Pasteurella multocida* (Walker & Weiss, 1994), indicating that these proteins consist of common structural building blocks that have been interchanged in the course of evolution. CNF1, the most thoroughly investigated representative of the CNF family, is a major virulence factor in uropathogenic *E. coli* (UPEC) strains, which live in the intestine and enter the urinary tract via the urethra (Boquet, 2001; Knust & Schmidt, 2010; Ho *et al*, 2018).

CNF1-containing strains exhibit a higher viability, have a higher potential to colonize the urinary tract, affect the function of immune cells and increase the inflammation rate (Falzano *et al*, 1993; Fournout *et al*, 2000; Rippere-Lampe *et al*, 2001). CNF1 was also identified in some intestinal and extraintestinal *E. coli* (ExPEC) where it was found to increase bacterial invasion into endothelial cells (Khan *et al*, 2002) and to promote malignant tumor conversion and intestinal cell invasiveness (Zhang *et al*, 2018; Fabbri *et al*, 2019). Similarly, the homologous toxin CNF_Y_, which shares 65% identity with *E. coli* CNF1, is crucial for the pathogenicity of *Y. pseudotuberculosis*, which causes food-borne and zoonotic enteric infections that manifest themselves as enteritis, mesenterial lymphadenitis and more rarely, in sequelae such as reactive arthritis (Koornhof *et al*, 1999a; Smego *et al*, 1999; Heine *et al*, 2018). The importance of CNF_Y_ is emphasized by the fact that a knock-out mutation of the *cnfY* gene leads to avirulence, allowing bacteria to become persistent in mice (Heine *et al*, 2018). Recent studies demonstrated that Rho GTPase activation by CNF_Y_ enhances the translocation of *Yersinia* outer proteins (Yops) into neutrophils and macrophages via a type III secretion system (T3SS). This blocks phagocytosis, triggers immune cell death and contributes to massive tissue damage by induction of pro-inflammatory responses and necrosis (Schweer *et al*, 2013; Wolters *et al*, 2013).

CNFs may also hold promise for treatment of neurological disorders and cancer (Maroccia *et al*, 2018). For example, CNF1 injection into brains of mice enhanced neuro-transmission and synaptic plasticity, leading to improved learning and memory (Diana *et al*, 2007). Moreover, CNF1 was able to rescue wildtype-like mitochondrial morphology in fibroblasts derived from patients with myoclonic epilepsy, and it reduced tumor growth (Vannini *et al*, 2016; Fabbri *et al*, 2018). As CNFs efficiently intoxicate a broad range of host cells, transport modules of the toxin may also be useful for drug delivery (Haywood *et al*, 2018). To exploit and further develop this tool, detailed knowledge of the molecular mechanisms underlying CNF secretion, translocation and activity is required. However, little is known about the global structure and the individual functional units of CNFs and so far, only the structure of the catalytic domain of CNF1 has been determined (Buetow *et al*, 2002).

At the sequence level, CNF-type toxins of different species share at least 55% overall identity (Fig EV1), indicating similar structures and conserved modes-of-action, although they show differential preferences with respect to the targeted Rho GTPase and interact with different host cell receptors (Hoffmann *et al*, 2004; Blumenthal *et al*, 2007). CNF1 uses two cellular receptors to enter host cells, the 37-kDa laminin receptor precursor p37LRP, which is recognized by sequences located within the N-terminus of the toxin, and the Lutheran adhesion glycoprotein/basal cell adhesion molecule (Lu/BCAM), which interacts with motifs in the C-terminal half (Fabbri *et al*, 1999; Chung *et al*, 2003; Kim *et al*, 2005; McNichol *et al*, 2007; Piteau *et al*, 2014; Reppin *et al*, 2017). The receptor of the N-terminal part of CNF_Y_ is still unknown, but it has been shown that binding of CNF1 to host cells has no effect on CNF_Y_ uptake (Blumenthal *et al*, 2007), suggesting that both toxins use different host cell factors to enter host cells. This has also been corroborated in a recent study that identified glycosaminoglycans as interaction partners of C-terminal fragments of CNF_Y_ (Kowarschik *et al*, 2020). The CNFs are taken up into endosomes and their release into the host cytoplasm requires two hydrophobic sequence motifs within the N-terminal half of the toxin that have been predicted to form α-helices. These helices are separated by a loop containing an acidic patch of four conserved acidic amino acids, and they are believed to insert into the endosomal membrane upon charge neutralization of the patch in the course of endosome acidification (Pei *et al*, 2001). An unidentified protease then cleaves CNF (i.e. CNF1 between residues 532 and 544), and the C-terminal fragment including the catalytic domain (residues 720-1014) is released into the cytosol of the host cell to mediate the cellular effects of the toxin (Pei *et al*, 2001; Knust *et al*, 2009).

In this study, we resolved the crystal structure of the full-length *Y. pseudotuberculosis* CNF_Y_ protein, necessary to achieve an understanding of its transport and functional mechanisms and its potential therapeutic use. The CNF_Y_ structure revealed a complex set-up of five individual building blocks and allowed us to obtain detailed information about the minimal secretion and translocation domain required to transport the catalytic domain or a fused cargo protein into the host cell cytosol, which could be exploited for drug delivery.

## Results

### CNF_Y_ contains five structural building blocks

Recombinant full-length CNF_Y_ was produced in *E. coli* (Appendix Fig S1) and crystallized in space group I2_1_2_1_2_1_. These crystals diffracted to 2.7 Å and contained one CNF_Y_ molecule in the asymmetric unit. Since no suitable search model for molecular replacement was available and crystallization of full-length seleno-*L*-methionine-labelled protein failed, we also crystallized different fragments of CNF_Y_: (i) one containing only the deamidase domain (residues 720-1014), (ii) another consisting of the subunit which is likely released into the cytosol (residues 526-1014) based on the homology to *E. coli* CNF1 (Hoffmann *et al*, 2004; Blumenthal *et al*, 2007), and (iii) a third fragment including the complete N-terminal portion with parts of the released subunit (residues 1-704) (Appendix Fig S1). A detailed description of structure determination by Se-SAD and molecular replacement is given in the Materials and Methods section and an overview over data collection and refinement statistics as well as the respective Protein Data Bank (Berman *et al*, 2000) deposition codes is provided in Table EV1.

CNF_Y_ adopts a compact, modular structure of five structural building blocks (D1-D5) with approximate dimensions of 115*73*65 Å (Fig 1A-C). All residues of the protein could be traced in the structure of the holo-protein with the exception of residues N430-K431, S550-L553 and P701-L717. The unresolved amino acids resided in surface loops, indicating intrinsic flexibility. Analysis with PiSQRD (Aleksiev *et al*, 2009) assigns domain boundaries to residues 1-22/135-424 (D1), 23-134 (D2), 425-529 (D3), 530-700 (D4) and 718-1014 (D5, deamidase domain). The compact arrangement of D1-D5 in the full-length structure of CNF_Y_ prompted us to investigate the interactions between the five individual domains of CNF_Y_ in more detail. Analysis with PISA (Krissinel & Henrick, 2007) reveals large hydrophobic interfaces between D1 and D2 (interface area 870 Å^2^) as well as between D3 and D4 (750 Å^2^) (Appendix Tab S1). The C-terminal domain D5 interacts mainly with D3 (610 Å^2^), which, as a consequence, partially blocks the entrance to the catalytic site. D5 interacts only weakly with D4 (380 Å^2^), which itself establishes an extensive interface with D1 (1390 Å^2^) (Fig 1C).

**Figure 1.**
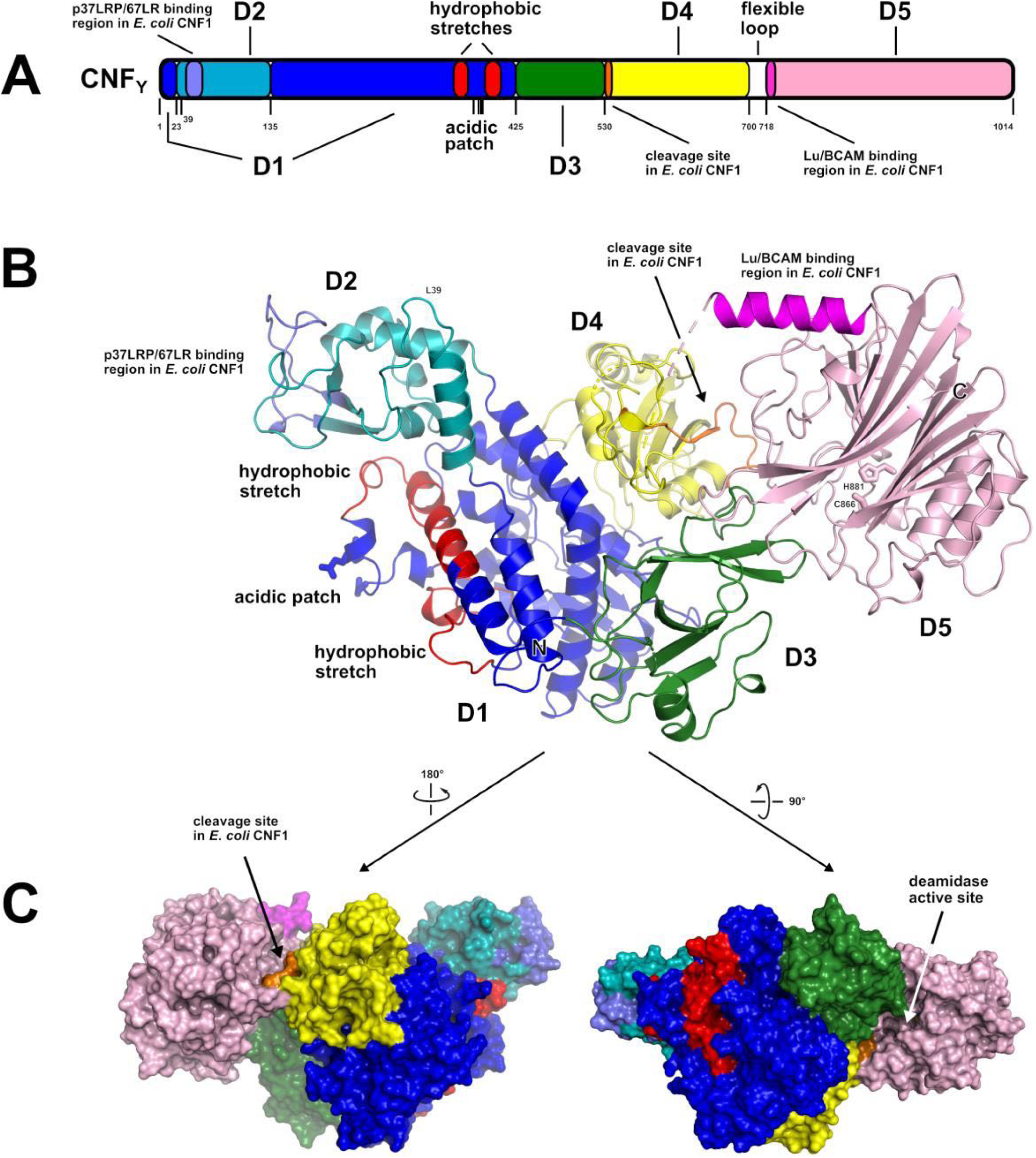
**The crystal structure of CNF_Y_ from *Y. pseudotuberculosis*.** A Domain boundaries and sequence motifs mapped to the sequence of CNF_Y_. B Cartoon representation of CNF_Y_, colored according to domain boundaries determined with PiSQRD (Aleksiev *et al*, 2009). Dark blue: domain D1, cyan: domain D2, dark green: domain D3, yellow: ADP-ribosyltransferase-like domain D4, pink: deamidase domain D5. Other colors indicate the position of sequence motifs that have been identified in *E. coli* CNF1, namely light blue: p37LRP/67LR receptor-binding motif, red: hydrophobic stretches predicted to form membrane-inserting α-helices, orange: cleavage site, magenta: main Lu/BCAM receptor-binding motif. The positions of N- and C-terminus are indicated by N and C, respectively. C Surface representation of CNF_Y_ as seen from two different orientations with respect to B. Note that the cleavage site between D3 and D4 (orange) as well as the deamidase active site in D5 are partially blocked in the structure of full-length CNF_Y_. The C-terminal domain D5 interacts mainly with D3 (610 Å^2^), which partially hides the catalytic site of D5, but it interacts only weakly with D4 (380 Å^2^), which itself establishes an extensive interface with D1 (1390 Å^2^) by mainly hydrophilic interactions (17 hydrogen bonds and 6 salt bridges).

The crystal structure of the fragment comprising residues 1-704 (D1-D4) is fully super-imposable with the respective residues of the holo-protein. The isolated D4-D5 fragment, on the other hand, showed a different orientation of the two domains with respect to the full-length protein and was thus included into the following more detailed structure-functional analysis together with full-length CNF_Y_.

### Structural analysis and structure-guided mutagenesis provide insight into the function of the structural building blocks of CNF_Y_

In order to gain insights into the biological function of the individual building blocks, we performed a detailed structural analysis of the full-length CNF_Y_ (D1-D5) and constructed truncated, mutated and marker-tagged versions of CNF_Y_ to investigate their secretion, translocation and enzymatic activity in human epithelial cells. The truncations were designed to interrupt the protein within linker regions between the individual domains (Fig 2A). CNF_Y_ variants were produced in *Y. pseudotuberculosis* or as recombinant proteins from *E. coli* (Fig 2B, Appendix Fig S1). Their enzymatic activity was tested with bacterial extracts or purified proteins by assessing their ability to deamidate RhoA in intact HEp-2 cells and host cell extracts, and by assessing the induction of actin rearrangements and the inhibition of cell division (formation of multinuclear cells) (Fig 3). The ability to bind to host cells was tested with 3xFlag-tagged versions of the CNF_Y_ derivatives (Figs 2D), and their capacity to reach late endosomes was measured with CNF_Y_-GFP fusion variants (Fig 4). CNF_Y_ export from the bacterial cell and cytosolic translocation of the CNF_Y_ activity domain were detected with CNF_Y_-β-lactamase (TEM) constructs (Figs 2C, 3A). Secretion of the CNF_Y_-TEM derivatives was analyzed by incubation of bacterial culture supernatants with nitrocefin, a chromogenic cephalosporin substrate used to detect β-lactamases. Translocation of the CNF_Y_-TEM fusion proteins was detected by staining the cytosol of host cells with the FRET substrate CCF4-AM, which contains the coumarin- and fluorescein-conjugated β-lactam cephalosporin and is green fluorescent (excitation at 409 nm, emission at 530 nm). Cleavage of the β-lactam ring shifts the fluorescence of the compound to blue (emission at 450 nm) and thus indicates the presence of TEM β-lactamase in the cytosol.

**Figure 2.**
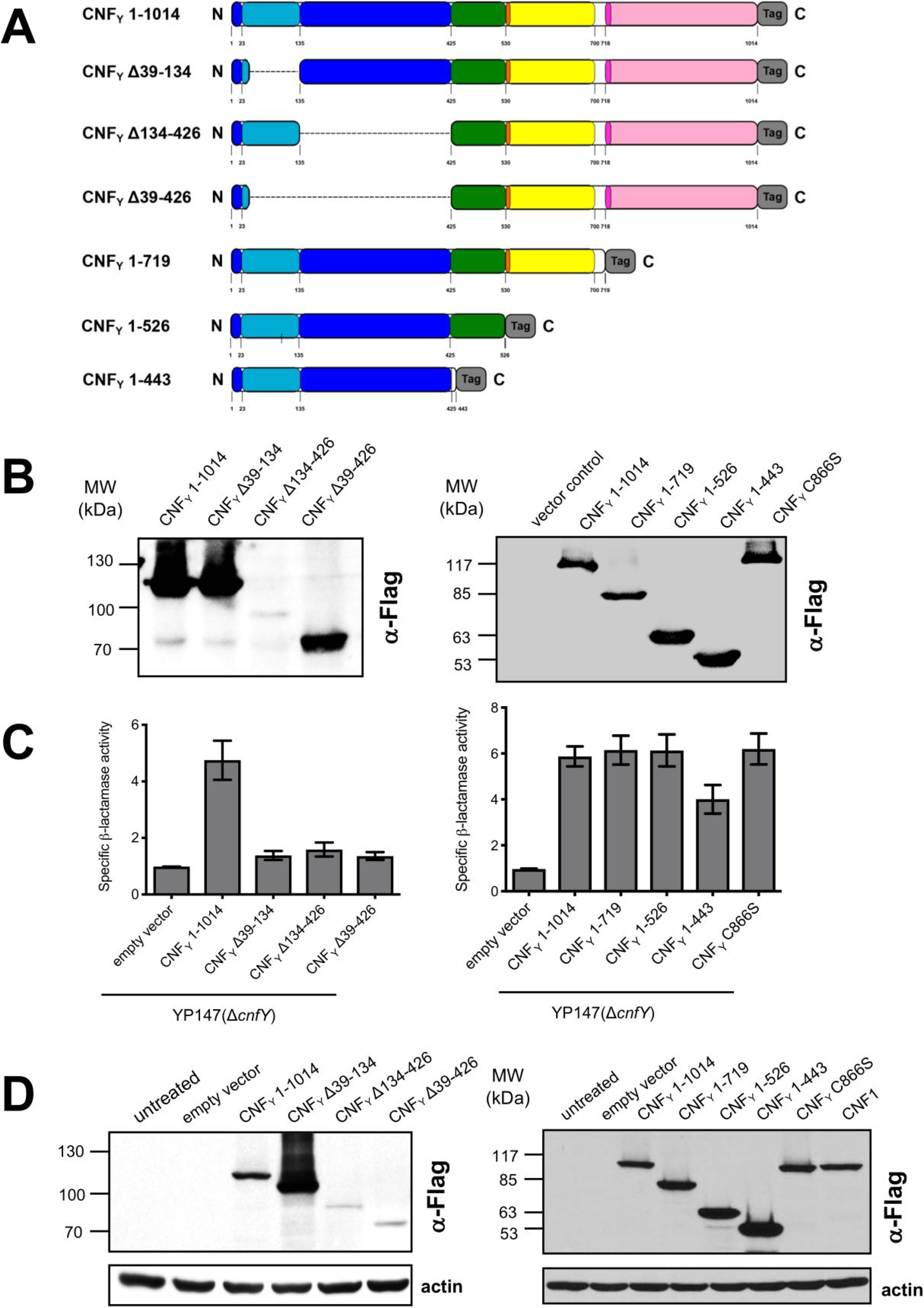
**Synthesis, secretion, and host cell binding of N- and C-terminal deletion variants of CNF_Y_.** A Schematic overview of marker-tagged CNF_Y_ deletion variants. B 3xFlag-tagged CNF_Y_ deletion variants were expressed in *Y. pseudotuberculosis* YP147 (Δ*cnfY*) from plasmids under control of their own promoter and were detected in whole cell extracts using an anti-Flag antibody. C To test secretion of the CNF_Y_ variants, full-length CNF_Y_ and different N- and C-terminally deleted variants fused to beta-lactamase (TEM) were expressed in *Y. pseudotuberculosis* YP147 (Δ*cnfY*). Beta-lactamase activity in the culture supernatant was subsequently measured using nitrocefin as substrate. D HEp-2 cells remained untreated or were incubated with 20 µg/ml of whole cell extract of *Y. pseudotuberculosis* expressing full-length CNF1, CNF_Y_ or the N- or C-terminally deleted toxin variants at 37°C for 4 h. The cells were thoroughly washed, pelleted, lysed and the toxin variants bound to the cells were identified by western blotting using an anti-Flag antibody.

**Figure 3.**
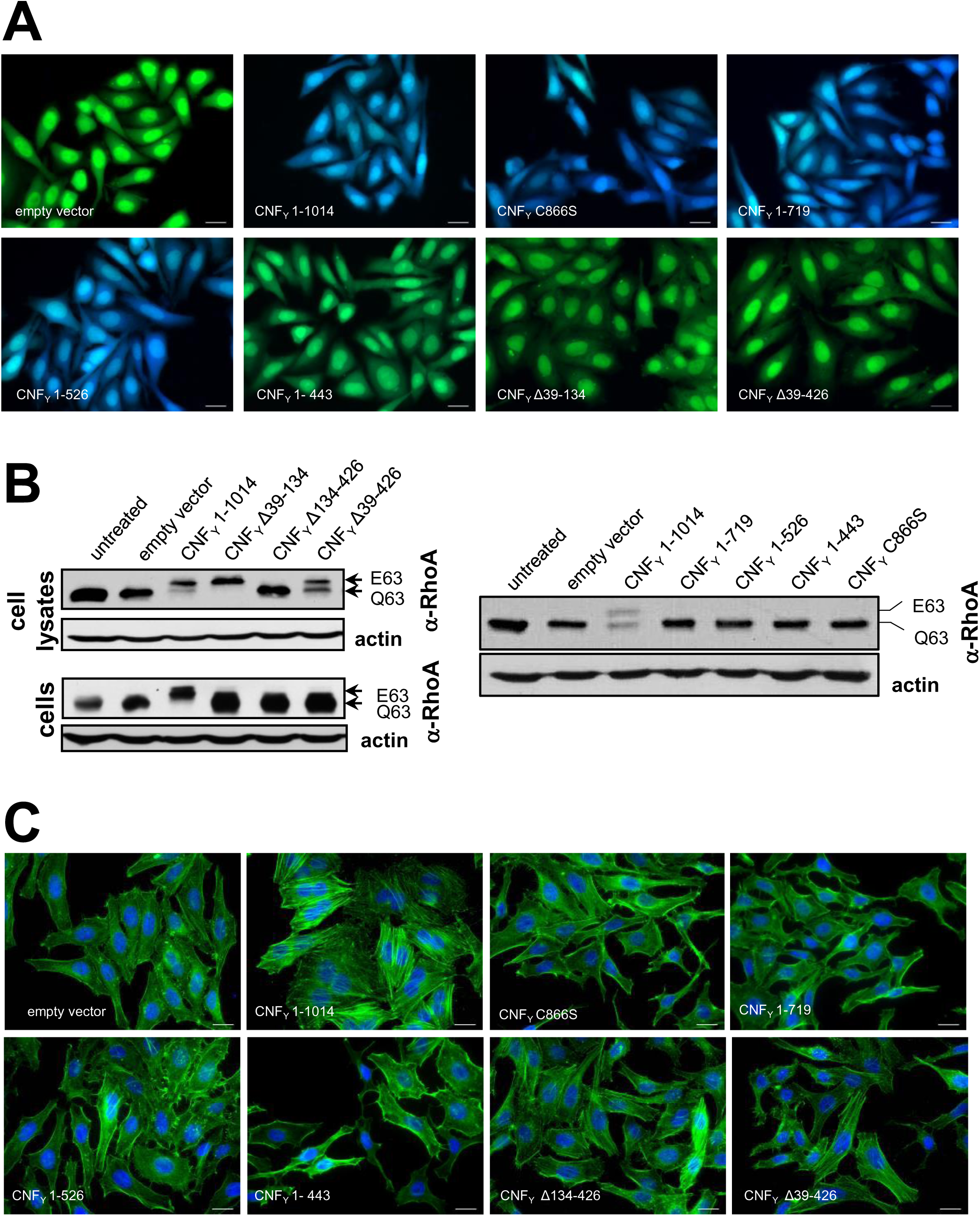
**Translocation of the deletion variants of CNF_Y_ and their influence on RhoA activation, actin rearrangements and multinucleation of host cells.** A HEp-2 cells were incubated with 20 µg/ml of whole cell extract of *Y. pseudotuberculosis* expressing full-length CNF_Y_ or the N- or C-terminally deleted toxin variants fused to β-lactamase (TEM) at 37°C for 4 h. Cleavage of the reporter dye CCF4-AM was used to visualize toxin delivery. After cell entry CCF4-AM is rapidly converted into the negatively charged form CCF4, which is retained in the cytosol and emits a green fluorescence signal (520 nm). In the presence of translocated β-lactamase fusion proteins, CCF4-AM is cleaved, and disruption of FRET results in blue fluorescence (447 nm). White bar: 20 µm. B Upper panels: HEp-2 cells remained untreated or were incubated with 20 µg/ml of whole cell extract of *Y. pseudotuberculosis* expressing full-length CNF_Y_ or the N- or C-terminally deleted toxin variants for 4 h. Cells were lysed and the deamidation of RhoA was analyzed by the shift of the modified Rho GTPase band in SDS PAGE gels; lower panel: HEp-2 cells were lysed and the cell extracts were incubated with full-length CNF_Y_ or the N-terminally deleted toxin variants for 4 h. The deamidation of RhoA in the cell extracts was analyzed by the mobility shift of the modified Rho GTPase on SDS PAGE after detection with anti-RhoA antibodies. C HEp-2 cells were incubated with 20 µg/ml of whole cell extract of *Y. pseudotuberculosis* expressing full-length CNF_Y_ or the N- or C-terminally deleted toxin variants for 24 h. The cell nuclei were strained with DAPI (blue) and the actin cytoskeleton was stained using FITC-phalloidin (green). The formation of large, multinuclear cells was observed by fluorescence microscopy and the formation of thick actin stress fibers and membrane actin folding were only observed with CNF_Y_-treated cells. The white scale bar is 40 µm. Cells incubated with extracts of YP147 (Δ*cnfY*) harboring the empty expression vector were used as negative controls.

**Figure 4.**
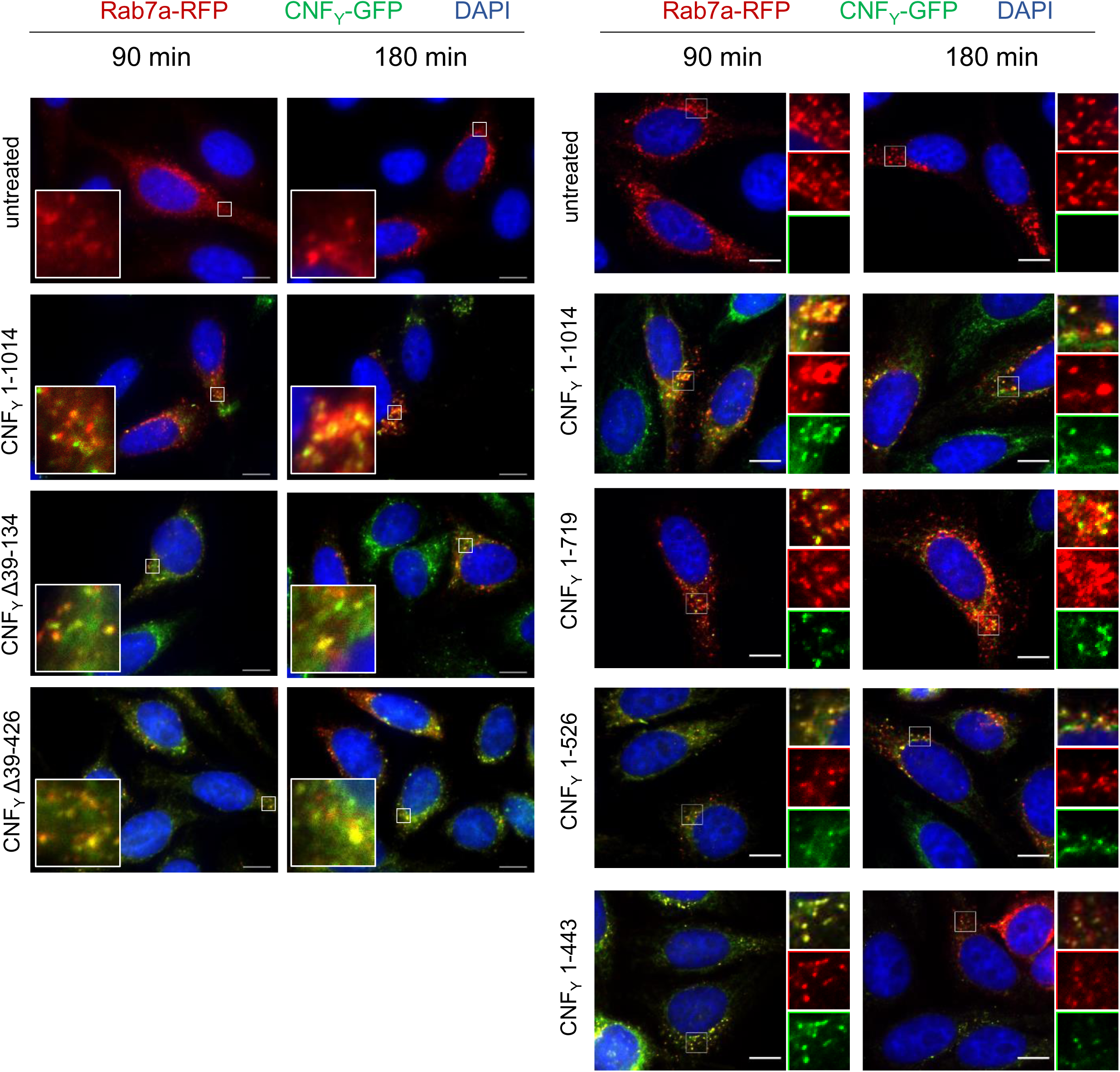
**Localization of the N- and C-terminal deletion variants of CNF_Y_ in the late endosome.** HEp-2 cells were incubated with 20 µg/ml of whole cell extract of *Y. pseudotuberculosis* expressing full-length CNF_Y_, N- or C-terminal deletion variants fused to GFP (green) for 90 or 180 min. Cells were fixed and processed for fluorescence microscopy. The red fluorescent signal represents late endosomes (CellLight Late Endosomes-RFP (Rab7a)). Nuclei were stained with DAPI (blue). A merged image of the different channels is shown, and smaller images are magnified views of boxed areas. White scale bar is 10 µm.

For validation, we first tested marker-tagged full-length wildtype CNF_Y_ (CNF_Y_ 1-1014) and a mutant derivative, namely CNF_Y_ C866S. This substitution inactivated the deamidase activity when introduced into *E. coli* CNF1 (Hoffmann *et al*, 2004). The marker-tagged versions of the full-length CNF_Y_ wildtype protein were efficiently produced (Fig. 2B), secreted (Fig 2C), and internalized into host cells (Fig 2D). Inside cells they were targeted to the late endosome (Fig 4) and translocated into the host cell cytosol (Fig 3A) to deamidate RhoA (Fig 3B) and induce formation of stress fibers or polynucleation in cultured cells (Fig 3C). In contrast, all CNF_Y_ C866S derivatives were found to abrogate the RhoA deamidation and induction of multinucleated cells, whereas other properties were not affected (Fig 2B-D, 3). Together, this demonstrates the suitability of the employed test systems.

#### Domain D1 is a major component of the translocation apparatus in CNFs

The crystal structure of CNF_Y_ revealed that domain D1 consists of separate areas covering residues 1-22 and 135-424, which together form a bundle of α-helices flanked by a four-stranded anti-parallel β-sheet that is covered with three α-helices from the other side (Fig 1B, Fig EV2). Sequences within domain D1 have previously been shown to contain elements that are required for translocation of the catalytic fragment of *E. coli* CNF1 (Pei *et al*, 2001; Knust *et al*, 2009), suggesting that this domain is a major component of the translocation machinery of CNFs.

Although domain D1 of CNF_Y_ is, due to its overall α-helical character, reminiscent of the translocation domain of other toxins such as diphtheria toxin (DT), searches with DALI (Holm & Rosenström, 2010) detected no significant structural homology to these proteins. Instead, it identified only the segment containing the four-stranded anti-parallel β-sheet (residues 152-343) as being somewhat similar to a fragment of the translocation domain of nigritoxin, a toxin of crustaceans and insects (PDB entry 5M41; 177 residues aligned, rmsd 3.8 Å, 11% sequence identity) (Fig 5A) (Labreuche *et al*, 2017). However, the translocation domain of nigritoxin is significantly smaller than the D1 domain of CNF_Y_ and does not contain hydrophobic sequence motifs that have previously been predicted to be essential for translocation in CNFs and other toxins (Pei *et al*, 2001; Orrell *et al*, 2017), hinting at distinct translocation mechanisms.

**Figure 5.**
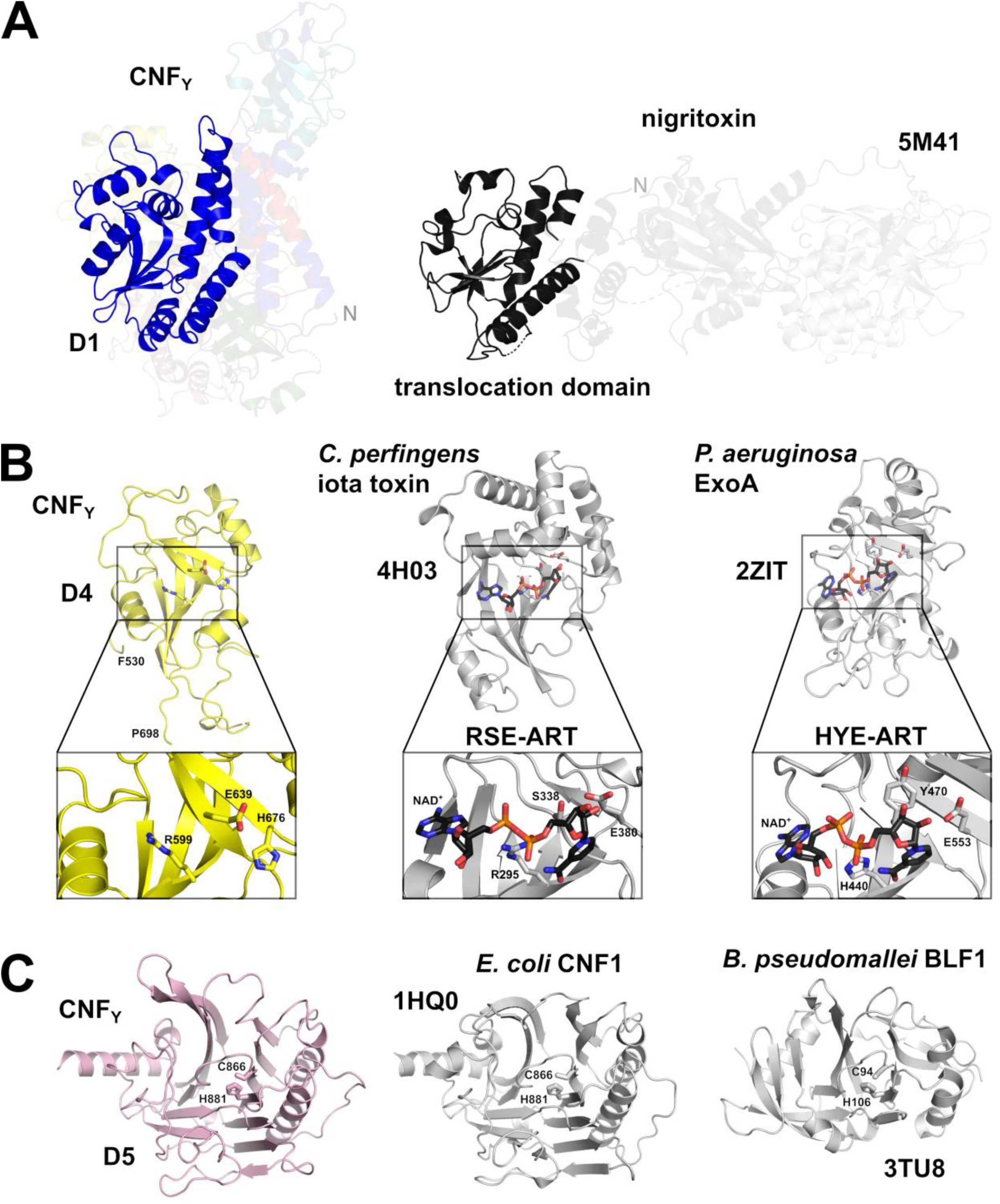
**Structural homology of the CNF_Y_ toxin and domain organization of toxins with a CNF-like translocation apparatus.** A Side-by-side comparison of CNF_Y_ and nigritoxin. Nigritoxin is a toxin of crustaceans and insects. The translocation domain of nigritoxin (PDB entry 5M41, (Labreuche *et al*, 2017)) and domain D1 of CNF_Y_ show partial structural similarity (highlighted areas). This similarity was identified with DALI (Holm & Rosenström, 2010) which was also used to align both structures. B The ART-like domain D4 of CNF_Y_. Essential residues of canonical ARTs are not conserved in CNF_Y_ (RSE-ARTs exemplified by *C. perfringens* iota toxin, PDB entry 4H03 (Tsurumura *et al*, 2013); HYE-ARTs exemplified by *P. aeruginosa* ExoA, PDB entry 2ZIT (Jørgensen *et al*, 2008); carbon atoms of NAD+ shown in black). C The deamidase domain D5 of CNF_Y_. C866 and H881 form a conserved catalytic dyad also found in the deamidase domain of *E. coli* CNF1 (PDB entry 1HQ0, (Buetow *et al*, 2001)) and *Burholderia pseudomallei* lethal factor BLF1 (PDB entry 3TU8, (Cruz-Migoni *et al*, 2011)).

To gain further insight into the function of D1, we constructed marker-tagged CNF_Y_ derivatives different deletions within the domain (Δ39-134, Δ134-426, Δ39-426; Fig 2A). The CNF_Y_ Δ134-426 derivative was less efficiently produced and inactive (i.e. it did not deamidate RhoA in host cell extracts, Fig 2B and 3B), indicating that it is improperly folded and less stable. However, the CNF_Y_ Δ39-134 and CNF_Y_ Δ39-426 proteins were well expressed and enzymatically active, yet, they failed to be secreted (Fig 2B-C), and consequently unable to trigger RhoA activation when added to host cells (Fig 3B). On the other hand, a C-terminally truncated CNF_Y_ protein containing the entire domain D1 (1-443) was secreted efficiently (Fig 2C), corroborating that this segment is necessary and sufficient for CNF_Y_ to exit the bacterial cell.

Use of CNF_Y_-TEM derivatives further revealed that all deletions of or within domain D1 abolished translocation of the active domain into the host cell cytoplasm when added to host cells (Fig 3A), although CNF_Y_ Δ39-134 and CNF_Y_ Δ39-426 derivatives were still able to bind and enter cells (Fig 2D), and associate with the late endosome (Fig 4). This demonstrated that domain D1 is an important component of the translocation machinery but not essential for host cell binding and endocytosis.

For *E. coli* CNF1, two hydrophobic α-helices have been predicted between residues 350-372 and 387-412, which are believed to insert into the endosomal membrane after charge neutralization of a conserved acidic patch in the connecting loop (D373, D379, E382 and E383) (Pei *et al*, 2001). However, the respective segments do not fold into the predicted α-helices in CNF_Y_ but adopt mostly loop-like structures with a helical part at their C-terminus (Fig 1B, Fig EV2), which may be a consequence of the neutral pH at which CNF_Y_ has been crystallized here. In line with previous work on *E. coli* CNF1, site-directed mutagenesis of the acidic residues E382 and E383 in the acidic patch to lysines did not abolish the enzymatic activity of the toxin derivative when added to cell lysates (Appendix Fig. S2). The CNF_Y_ E382K/E383K variant was still secreted and able to enter cells but failed to activate RhoA and induce multinucleation (Appendix Fig S2). This supported previous assumptions that acidic residues in the connecting loop are important for translocation (Pei *et al*, 2001). Importantly, domain D1 alone does not seem to be sufficient for the translocation process, as the C-terminally truncated derivative CNF_Y_ 1-443, consisting of domains D1-D2, was unable to translocate the β-lactamase TEM cargo protein into the cytosol, deamidate RhoA in host cells and induce multinucleation (Fig 3), despite the fact that it was internalized and reached late endosomes (Fig 4). This indicated that the translocation machinery of CNFs requires additional components for functionality.

#### Domain D2 is a receptor binding domain of CNFs

Unlike the N-terminal α-helix (residues 5-18), which is an integral part of the helical bundle that dominates domain D1, residues 23-134 seem to establish a separate structural building block (domain D2) that protrudes from the mostly α-helical subunit, potentially suggesting its insertion during evolution (Fig 1B). It consists of a three-stranded anti-parallel β-sheet flanked by α-helices at the side facing D1 and by several surface-exposed loops at the other. One solvent-exposed loop contains residues 53-75, a segment that has previously been implicated in host cell binding of *E. coli* CNF1 to receptor p37LRP/LR67 (Fig 1B, Fig EV2) (Fabbri *et al*, 1999; Chung *et al*, 2003; Kim *et al*, 2005). It is hence conceivable that this domain represents the N-terminal receptor binding domain of CNFs. The solvent-exposed loop interacts with residues 112 to 116 at the surface of D2, such that these segments together may establish a receptor-binding interface of the CNFs (Fig EV2). There are ten amino acid changes between CNF1 and CNF_Y_ within these regions (Fig EV1), which may account for the distinct receptor specificity of both toxins (Blumenthal *et al*, 2007).

As outlined above, host cell binding and colocalization studies with truncated CNF_Y_ derivatives showed that the D1-2 (1-443) fragment was able to bind host cells (Fig 2D) and reached the late endosomes (Fig 4), indicating that this truncated version of CNF_Y_ indeed includes a cell receptor binding site similar to CNF1 (Fabbri *et al*, 1999; Chung *et al*, 2003; Kim *et al*, 2005). However, this also demonstrates that CNF_Y_, unlike previous work with CNF1 suggests (Piteau *et al*, 2014; Reppin *et al*, 2017), does not require a second recognition site in the C-terminal region to enter host cells.

#### Domain D3 is an essential part of the translocation apparatus of CNFs

The third domain, D3 (residues 425-529), is reminiscent of an incomplete β-barrel, containing six anti-parallel strands in CNF_Y_. No homologous structures could be discovered with DALI. Because a fragment consisting only of domains D1-2 (CNF_Y_ 1-443) was not able to translocate the fused TEM-β-lactamase into the host cell cytosol whereas the D1-3 fragment (CNF_Y_ 1-526) was (Fig 3A), D3 is obviously essential for intoxication. D1-3 translocated TEM-β-lactamase with the same efficiency as full-length CNF_Y_ (Appendix Fig. S3), and although it is unclear whether β-lactamase was released from D1-3 or only exposed on the cytosolic side from the endosomal membrane, these experiments clearly demonstrated that the translocation apparatus of CNFs consists of the three domains D1-3. The importance of D3 is also highlighted by the fact that (i) the N-terminus of *Pasteurella multocida* exotoxin PMT possesses high homology to D1-3 of CNF_Y_, whereby cargo delivery includes proteolytic cleavage downstream of the prospective D3 domain, and (ii) the finding that a hybrid toxin consisting of this N-terminal fragment (residues 1-505) of PMT and the ADP-ribosylating domain of DT was able to intoxicate cells (Bergmann *et al*, 2013; Clemons *et al*, 2018).

#### The cleavage site between D3 and D4 is partially shielded in full-length CNF_Y_

The imperfect β-barrel domain D3 and the following domain D4 are connected via a linker that is partially shielded by the C-terminal deamidase domain D5 in the structure of full-length CNF_Y_ (Fig 1C). In CNF1, this linker is cleaved between residues 532-544 to release the D4-5 subunit into the host cell cytosol (Knust *et al*, 2009), suggesting that the respective segment must become solvent-exposed in the course of host cell intoxication to be accessible to proteases. As outlined below, this likely is linked to unfolding during translocation of the D4-5 segment into the cytosol of the host cell.

In order to identify amino acids that are important for the cleavage step, we introduced mutations within this linker. All recombinant mutant proteins were able to bind to host cells and deamidate RhoA in cell lysates, indicating proper folding and full activity (Fig 6). Interestingly, the CNF_Y_ I535L/P536A/V537G mutant with changes in the N-terminal part of the linker (CNF_Y_ mut1) promoted translocation into the host cell cytoplasm and was able to deamidate RhoA when added to host cells (Fig 6). However, this was not the case for the CNF_Y_ variant I535L/P536A/V537G/F539L/D541A/K542A (Fig 6), suggesting that this variant is unable to escape the endosome. In summary, this indicated that amino acids important for proper processing are located at the C-terminal end of the linker.

**Figure 6.**
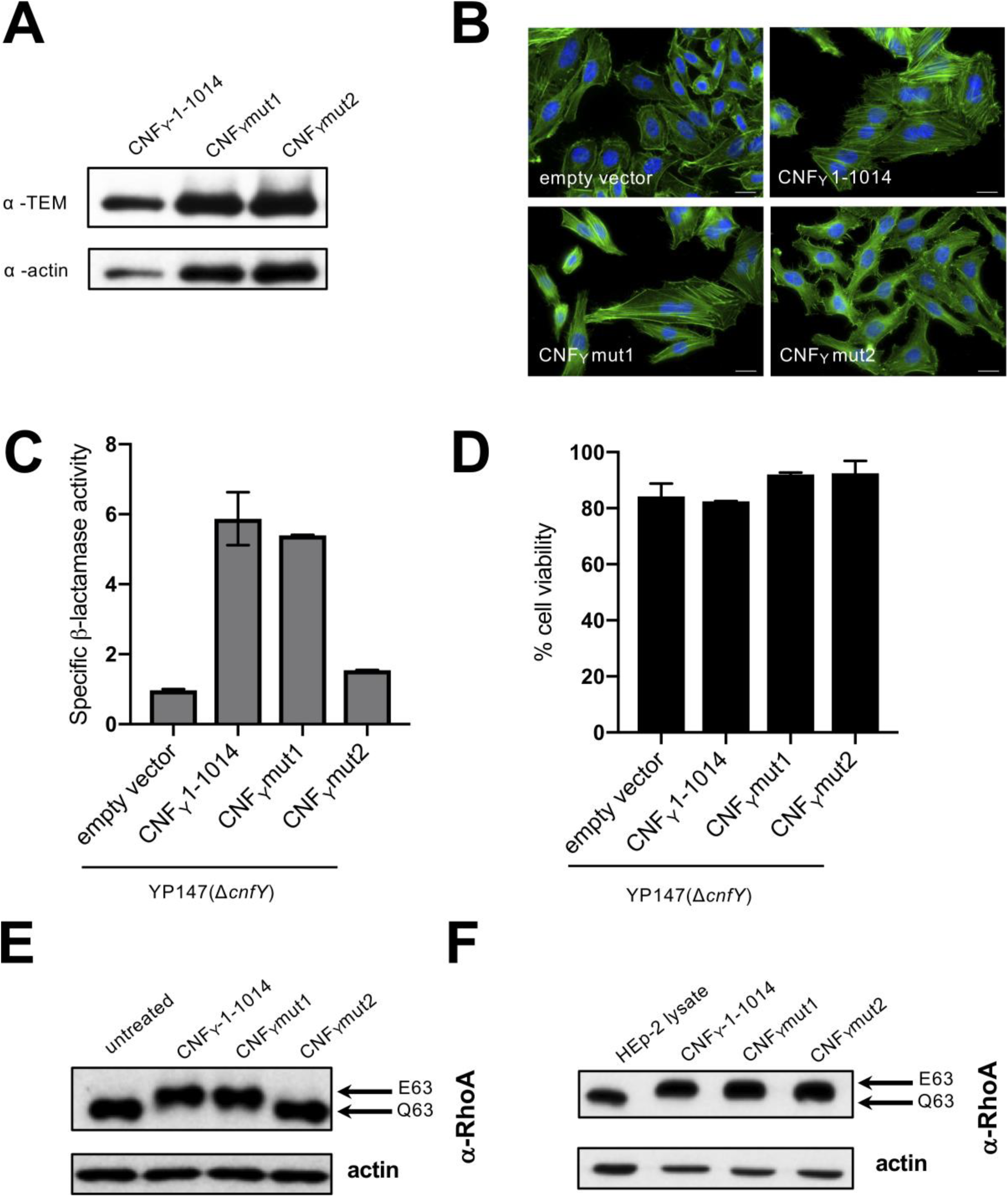
**Characterization of mutants in the linker region connecting domain D3 and D4.** A HEp-2 cells were incubated with 20 µg/ml of whole cell extract of *Y. pseudotuberculosis* expressing CNF_Y_, the toxin variant mut1: CNF_Y_I535L/P536A/V537G or mut2: CNF_Y_I535L/P536A/V537G/F539L/D541A/K542A fused to TEM or no CNF_Y_ protein (empty vector) for 4 h. Cells were lysed and the binding of the different CNF_Y_ proteins to HEp-2 cells was analyzed by immunoblotting. B HEp-2 cells were incubated with 20 µg/ml of whole cell extract of *Y. pseudotuberculosis* expressing CNF_Y_, the toxin variant mut1: CNF_Y_I535L/P536A/V537G or mut2: CNF_Y_I535L/P536A/V537G/F539L/D541A/K542A or no CNF_Y_ protein (empty vector). The cell nuclei were stained with DAPI (blue) and the actin cytoskeleton was stained using FITC- phalloidin (green). The results indicated the formation of polynucleated cells and stress fibers only in cells treated with CNF_Y_ and CNF_Y_I535L/P536/V537G. The white scale bar is 20 µm. C Nitrocefin (2 mM) was added to the supernatant from 25°C overnight *Yersinia* cultures expressing the indicated CNF_Y_ derivatives to determine β-lactamase activity by measuring changes in absorbance at 390 nm (yellow) and 486 nm (red). D The viability of *Y. pseudotuberculosis* YPIII expressing the indicated CNF_Y_ derivatives was assessed in equalized bacterial cultures using the BacTiter-Glo Microbial Cell Viability Assay kit (Promega). E HEp-2 cells treated with 20 µg/ml of whole cell extract of *Y. pseudotuberculosis* expressing indicated CNF_Y_ variants for 4 h were lysed and the deamidation of RhoA was analyzed by the mobility shift of the modified RhoA GTPase detected by immunoblotting. F The activity of the CNF_Y_ derivatives was tested by analyzing the deamidation of RhoA in HEp-2 cell lysates by the mobility shift of the modified GTPase detected by immunoblotting.

#### Domain D4 has structural similarity to ADP-ribosyl transferases

Sequence analysis places the fourth domain D4 (residues 530-700) into the DUF4765 family, a building block that is also found in a number of other uncharacterized bacterial proteins (Fig EV3A). Surprisingly, structure similarity searches reveal distant but significant homology to ADP-ribosyl transferase (ART) domains (Appendix Tab S2), which are wide-spread in protein toxins (Fieldhouse & Merrill, 2008). This similarity is exemplified by two examples (*Clostridium perfringens* iota toxin, PDB entry 4H03 (Tsurumura *et al*, 2013); *Pseudomonas aeruginosa* ExoA, PDB entry 2ZIT (Jørgensen *et al*, 2008)) shown in Fig 5B and Fig EV3.

In order to gain further insight into the function of domain D4, which is translocated and released together with the catalytic domain, we analyzed two mutant variants harboring internal deletions of amino acids 527-719 or 527-699. However, both variants did not deamidate RhoA in host cell lysates, indicating that these mutant proteins failed to fold properly (Appendix Fig S4). Since structure similarity searches revealed distant homology to the ADP-ribosyltransferase (ART) domains, we hypothesized that CNFs may possess a second, previously unrecognized enzymatic function encoded in D4. The active sites of ARTs fall into two groups, the RSE- for a conserved arginine-serine-glutamate active site motif and the HYE-ARTs for histidine-tyrosine-glutamate (Cohen & Chang, 2018). The ART-like domain D4 contains arginine, glutamate and histidine at the respective positions instead (R599, E639, H676), which could, in principle, support similar chemistry and are, with the exception of the CNF-specific H676, highly conserved in other DUF4765-containing proteins (Fig EV3A). However, exchange of CNF_Y_ E639, which is in the comparable position of the conserved glutamate of bacterial ARTs, to alanine or glutamine had no effect on CNF_Y_ function (Appendix Fig S5). Moreover, we could not detect binding of the ART cosubstrate NAD^+^ to a CNF_Y_ fragment consisting of domains D4-5 in microscalar thermophoresis titration experiments. The inability to bind NAD^+^ may be a consequence of the altered geometry of the potential NAD^+^ binding site of D4, which leads to a shallower hypothetical NAD^+^ binding site and to clashes when NAD^+^ is superimposed from the two examples mentioned above (Fig EV3B).

#### Domain D5 is highly similar to the deamidase domain of CNF1

The C-terminal catalytic deamidase domain D5 is linked to D4 via the unstructured residues P701-L717 (Fig 1B). These belong to the postulated binding epitope for the Lu/BCAM host receptor of CNF1 (Piteau *et al*, 2014), and their flexibility may be a requirement for receptor binding by the toxin.

The D5 domain is very similar to the respective domain of *E. coli* CNF1 (PDB entry 1HQ0 (Buetow *et al*, 2001); 1.8 Å rmsd over 295 residues, 59% sequence identity), featuring a central β-sandwich with shielding α-helices on both sides (Figs 1B and 5C). The active site employs the conserved cysteine/histidine couple C866/H881 (Hoffmann *et al*, 2004), which lies in a crevice on the surface of the domain and deamidates a conserved glutamine in the switch-II region of the targeted Rho GTPases. In agreement, only the full length CNF_Y_ protein was able to activate the Rho GTPase RhoA (Fig 3B), resulting in the induction of polynucleation in living cells (Fig 3C), whereas all protein variants with deletions of the C-terminal domain D5 eliminated toxicity (Figs 3B-C). This is consistent with studies showing that the C-terminal 300 amino acids (709-1014) of the related *E. coli* CNF1 protein are important for its activity (Koornhof *et al*, 1999b; Zhang *et al*, 2018; Fabbri *et al*, 2019). Neither the deletion nor site-directed mutagenesis of the catalytic domain affected secretion, host-cell binding, or protein translocation (Figs 2B-C and 3A), indicating that the sole role of the C-terminal domain is the targeting and modification of Rho GTPases.

While the crevice is nearly identical between CNF_Y_ and CNF1, differences exist between two loops at the periphery of the active site, namely residues 961-970 and residues 996-1004 (Fig 5C, Fig EV1). It is not clear if these deviations are sufficient to explain the slightly altered preferences for various Rho GTPases that have been described for both CNFs (Hoffmann *et al*, 2004; Schweer *et al*, 2013), but amino acids in one of these loops (residues 961-970) has previously been implicated in substrate recognition of CNF_Y_ and CNF1 (Hoffmann *et al*, 2007). Interestingly, a similar deamidation domain is also found in *Burkholderia* lethal factor 1 (BLF1), despite extremely low sequence similarity (Fig 5C; PDB entry 3TU8; (Cruz-Migoni *et al*, 2011); 3.5 Å rmsd over 169 residues, 7% sequence identity). BLF1 is a single-domain toxin that deamidates a glutamine residue in translation initiation factor eIF4A. In addition, several other toxins including e.g. *Bordetella pertussis* dermonecrotizing toxin DNT are predicted to contain similar deamidase domains (Ho *et al*, 2018).

#### The D4-5 fragment of CNF_Y_ contains a receptor binding site sufficient for endosomal uptake

We further characterized the properties of the recombinant D4-5 domains (residues 526-1014), which together constitute the C-terminal fragment that is translocated into the host cell cytoplasm after cleavage of CNF_Y_ (Fig EV4). Although catalytically active when added to host cell extracts, D4-5 is unable to deamidate RhoA when added to host cells (Fig EV4E-D). This suggested that it was either not taken up into the host cell or unable to escape the endosome to reach the cytoplasm. Cell binding assays demonstrated, however, that the D4-5 fragment (CNF_Y_ 526-1014), similar to D1-3 (CNF_Y_ 1-526), specifically interacts with host cells (Fig 2D, Fig EV4C and S6). A parallel analysis of D3-5 (CNF_Y_ 426-1014), also missing the N-terminal receptor binding domain, confirmed these results (Fig EV4C). This indicated the presence of a second host cell binding site in the C-terminal region, which is strongly supported by the fact that N-terminal deletions missing parts of D1-2 (CNF_Y_ Δ39-426) are still able to promote cell binding and endosomal uptake as indicated by colocalization studies (Figs 2D and 4).

In *E. coli* CNF1, the binding site for the Lu/BCAM receptor was shown to include amino acids 720-730 of the catalytic domain (Reppin *et al*, 2017). The respective segment is significantly different in CNF_Y_ (Fig EV1), which may explain why CNF_Y_ does not interact with Lu/BCAM. Instead, a recent study has shown that a C-terminal fragment of CNF_Y_ (residues 709-1014) employs glycosaminoglycans as receptors and is sufficient for endosomal uptake (Kowarschik *et al*, 2020), in line with the observations made here. Thus CNF_Y_, similar to CNF1 (Piteau *et al*, 2014; Reppin *et al*, 2017), contains two distinct host cell binding sites, one each at the N- and C-terminus. In CNF_Y_, these binding sites enable endosomal uptake independently of each other, and the presence of two receptor binding sites might broaden the range of targeted cells or may increase host cell binding affinity.

#### The D4-5 segment adopts a different conformation and shows increased deamidase activity after cleavage from full-length CNF_Y_

Previous work with *E. coli* CNF1 demonstrated that translocation releases a fragment consisting of the ART-like domain D4 and the deamidase domain D5 into the cytosol of host cells (Knust *et al*, 2009). This prompted us to crystallize the respective segment of CNF_Y_, leading to a structure in which linker residues V702-L717 again were too flexible to be traced and in which the two domains adopt a different relative orientation with respect to the full-length protein. Whereas D4 and D5 interacted only weakly in the complete toxin, they now engage in a large interface (1100 Å^2^), whereby the active site crevice of D5 is extended by D4 and becomes fully solvent-exposed (Fig 7A). To reach this position, domain D4 has to rotate by more than 140°, which can probably only be achieved after cleavage from D1-3 and through the flexibility of the linker connecting both domains. The contact area between both domains in the free D4-5 subunit overlaps largely with that of D3-5 in the full-length structures such that both conformations are mutually exclusive, i.e. the D4-5 subunit cannot adopt the conformation observed in the free state when it is bound to D1-3.

**Figure 7.**
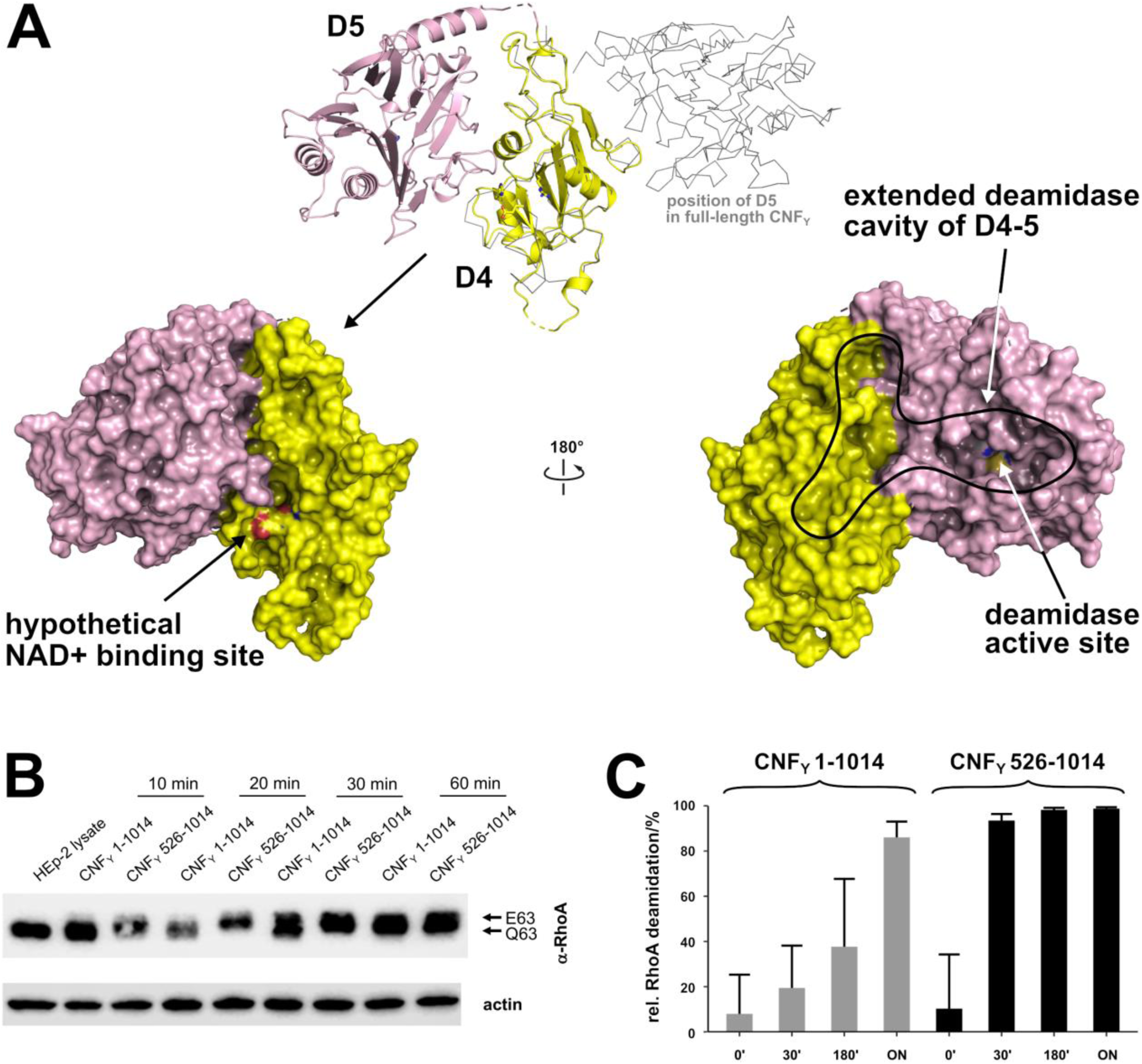
**Structure and deamidation activity of the free D4-5 subunit of CNF_Y_.** A Crystal structure of the free D4-5 subunit. Note the different relative orientations of domains D4 and D5 with respect to the structure of full-length CNF_Y_ (top, thin grey lines). The domain D4 forms a large interface area (1100 Å^2^) with the catalytic domain D5 involving several polar interactions (8 hydrogen bonds and 8 salt bridges), whereby the active crevice is extended and fully solvent-exposed as can be seen in the right surface plot at the bottom of the panel. The hypothetical NAD^+^ binding site of the ART-like D4 domain is located on the opposite face (left surface plot). Note that the deamidase active site of domain D5, unlike in the full-length structure (Fig 1), is fully accessible and that its extended shape is also determined by domain D4. B Comparative analysis of RhoA activation in HEp-2 cell lysate by CNF_Y_ and the recombinant D4-5 protein. Purified CNF_Y_ or the D4-5 fragment (1 µM) was added to extracts of HEp-2 cells and incubated for 10, 20, 30 or 60 min. Deamidation of RhoA was analyzed by the shift of the modified Rho GTPase band in SDS PAGE gels after detection with anti-RhoA antibodies. C Comparative analysis of recombinant RhoA deamidation by CNF_Y_ and the D4-5 fragment. Recombinant RhoA was incubated with purified CNF_Y_ or the D4-5 fragment and samples were separated by SDS-PAGE after the indicated times before subjecting to trypsin digestion and quantification of deamidation of Q63 by mass spectrometry. Error bars represent standard deviations of triplicate measurements.

The finding that the active site within domain D5 becomes more exposed in the recombinant D4-5 fragment suggested that presence of domains D1-3 may repress the deamidase activity of CNF_Y_ and that cleavage of the D4-5 fragment is required to rearrange and liberate and activate the catalytic unit. To test this hypothesis, we incubated equal amounts of purified CNF_Y_ 1-1014 (D1-5) and CNF_Y_ 526-1014 (D4-5) with host cell extracts or recombinant RhoA and analyzed the deamidation of RhoA in gel-shift assays and with proteomic methods. As shown in Fig 7B-C, RhoA deamidation by CNF_Y_ 526-1014 (D4-5) protein was indeed significantly faster than by the full-length protein, indicating that the domain rearrangement seen in the D4-5 fragment enhances the deamidase activity and that the full-length toxin is in an autoinhibited state with respect to this activity.

## Discussion

Here we show that CNF_Y_ and related CNFs consist of five individual structural building blocks that enable the different steps of the intoxication process, namely secretion, cell attachment, entry, translocation and enzymatic activity. The three N-terminal domains D1-3 all possess novel folds and constitute the secretion and membrane translocation unit, whereas the two C-terminal domains D4-5 form the toxicity-mediating unit of the toxin. We further show that the D1-3 unit is sufficient to transport cargo proteins such as β-lactamase into the cytosol of host cells. Strikingly, both the Rho deamidation and β-lactamase activity were preserved when the reporter was fused to the C-terminal end of the full-length protein (Fig 3), indicating that the secretion and transport module of the CNF_Y_ protein is very robust and insensitive to C-terminal extensions, making it an attractive tool for drug delivery.

The molecular mechanisms by which domains D1-3 promote the delivery of the cargo from the bacterial to the host cell cytosol are still unknown. However, considering the compact arrangements of the modules with large hydrophobic interfaces between domains D1 and D2, D3 and D4, as well as D3 and D5, it is likely that the full-length toxin is secreted from the bacterial cell and endocytosed by the host cells as monolithic compact structure. In fact, the CNF_Y_ toxin has recently been identified on the surface of outer membrane vesicles (OMVs) isolated from *Y. pseudotuberculosis* culture supernatants (Monappa et al. 2018). While this could indicate that the toxin might be predominantly delivered into endosomes by OMVs, our data further show that also the purified CNF_Y_ toxin interacts with and is efficiently internalized into host cells on its own. This suggests that the toxin is also directly secreted by the bacterial cell and/or exposed on the OMVs to promote contact with target cells.

The data presented here further show that the cellular toxin uptake process not only requires segments identified for receptor binding in D2 and for translocation in D1, but also the imperfect β-barrel domain D3. It is interesting to note that D1 is, due to its mostly α-helical character, reminiscent of the translocation machinery of other toxins including that of the diphtheria toxin DT. DT, CNFs and several other AB-type toxins contain two hydrophobic stretches that are believed to fold into α-helices and insert into the endosomal membrane after charge neutralization of surrounding acidic residues (Pei *et al*, 2001; Orrell *et al*, 2017). In CNF_Y_, the respective residues 350 – 372 and 387 – 412 were not found in the predicted α-helical structure (Fig 1B). However, since the crystal structures presented here have been obtained at neutral pH, it is conceivable that this region undergoes refolding during endosomal acidification. While the precise molecular mechanism of the complex translocation process of CNF_Y_ is still unclear (Pitard & Malliavin, 2019), work with DT suggests that the catalytic subunit of this toxin is unfolded in the translocation process (Murphy, 2011). The fact that (i) translocation in CNFs also involves two hydrophobic motifs interrupted by acidic residues and (ii) the observation that a sequence that gets cleaved to release the catalytic unit (D4-5) of CNF (residues 532 to 544) is not accessible in the full-length structure (Fig 1C) may hint at a similar unfolding in the CNFs. In this respect, the similarity of parts of the CNF_Y_’s D1 domain to the putative translocation domain of nigritroxin (Fig 5A) (Labreuche *et al*, 2017) is interesting, because the translocation domain of this toxin does not contain hydrophobic α-helices. This could indicate that the translocation process occurs through several steps that involve different parts of the translocation machinery, most of which are not shared between the CNFs and nigritoxin.

Sequence searches in the UniRef50 database (Suzek *et al*, 2015) revealed that large sections of the D1-3 domain of CNF_Y_ are also found in a number of un- or less characterized bacterial proteins, suggesting that these proteins are toxins that apparently utilize a similar secretion and translocation device for their catalytic domains (Fig 8). For example, about 530 amino acids of the N-terminus of CNF_Y_ share between 30-50% sequence identity to the N-terminus of *Pasteurella multocida* toxin PMT (Bergmann *et al*, 2013). Moreover, members of a group consisting of 372 proteins with approximately 900 residues each were found to possess a canonical RSE-type ART-domain at their C-terminus (represented by UniProt entry A0A0P9UH04 from *Pseudomonas syringae pv. maculicola*) in addition to a CNF-like translocation apparatus. A second group of 206 proteins with more than 1000 residues contains a C-terminal glycosyltransferase (represented by UniProt entry A0A0N8SZE6 from *Pseudomonas syringae pv. syringae*). Since the C-termini of these proteins differ from CNF_Y_ (Fig 8), it is likely that the toxins consist of individual modules that have been shuffled in the course of evolution. This aligns with a recent analysis of the distribution of CNF-like deamidase domains, which are also found at different positions within the sequence of other toxins or as stand-alone proteins (Cruz-Migoni *et al*, 2011; Ho *et al*, 2018).

**Figure 8.**
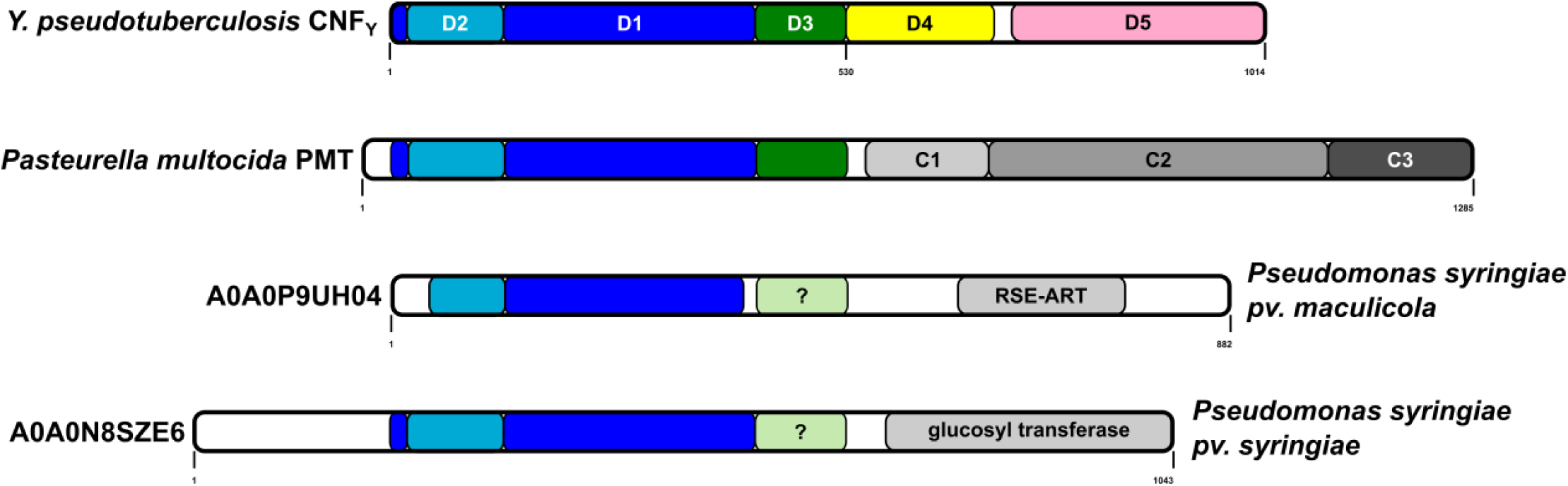
**Architecture of bacterial toxins with a CNF-like translocation apparatus** Shown are the architecture of CNF_Y_, *Pasteurella multocida* toxin PMT and two uncharacterized proteins from *Pseudomonas syringiae*. The released fragment of PMT contains three domains of which C1 is required for membrane binding, the C2 domain has an unknown function and the C3-domain activates heterotrimeric G-proteins by deamidation. The two *Pseudomonas syringiae* proteins A0A0P9UH04 and A0A0N8SZE6 represent two uncharacterized toxins that encode catalytic domains of the indicated type. While sequence alignments unequivocally reveal a CNF-like imperfect β-barrel in PMT, the presence of this domain in the *P. syringiae* toxins is less obvious.

The finding that the DUF4765 domain D4 shows similarity to ADP-ribosyltransferases was surprising and led us to investigate if the released D4-5 may possess an additional and previously unrecognized enzyme activity that may contribute to the toxicity of CNFs. However, the observation that CNF_Y_ toxicity strictly depended on the activity of the deamidase D5 domain whereas mutations of the potential/suggested NAD^+^ binding domain within D4 had no effect (Appendix Fig S5), together with the fact that we could not detect NAD^+^ binding in biophysical experiments, speaks against such an additional activity. On the other hand, the NAD^+^ affinity of some ADP-ribosyl transferases is very low. This is exemplified by cholera toxin, where a K_D_ of 4.0 ± 0.4 mM has been determined (Galloway & van Heyningen, 1987). Alternatively, the grossly different and mutually exclusive relative orientations of the D4 and D5 domains in the free D4-5 subunit with respect to the full-length CNF_Y_ structure (Fig 7) could suggest that D4 may have a regulatory role. On the one hand, the finding that the active site becomes solvent-accessible and that the crevice leading to the active site of D5 becomes extended by parts of D4 (Fig 7) could fine-tune the deamidase function of D5 with respect to general activity levels or substrate specificity towards RhoA, Rac1 or Cdc42. On the other hand, D4 could contribute to localizing the catalytical unit within the host cell by promoting access to membrane-associated Rho GTPases. In fact, CNFs act predominantly on Rho GTPases bound to GTP, a form essentially found at the cytoplasmic face of the host cell membrane (Boquet, 2001). Clearly, the importance of D4 merits future studies.

In summary, the data presented here provide insight into the full-length and released active D4-5 structure, and they illustrate the importance of the individual building blocks of CNFs and related exotoxins. This not only forms the basis for the detailed analysis of the molecular secretion and transport mechanism, but also enables the rational design of the transport module as a toxin-based cargo delivery tool for cytosolic drug/therapeutics delivery and the structure-guided development of inhibitors of CNF-like virulence factors.

## Materials and Methods

### Bacterial strains, cell lines, plasmids and growth conditions

All bacterial strains and plasmids used in this study are listed in Appendix Tab S3. All oligonucleotide primers used for cloning are listed in Appendix Tab S4. *E. coli* strains were grown in Luria-Bertani (LB; Becton Dickinson) broth at 37°C. *Yersinia* strains were aerobically grown in LB at 25°C or 37°C. Other media used for bacterial growth were brain-heart infusion broth (BHI) (Gibco) and Double Yeast Tryptone medium (DYT) (Gibco). Cultures were supplemented with 30 μg/ml kanamycin (Kan) or chloramphenicol (Cm) where necessary. HEp-2 cells (ATCC CCL-23) were grown at 37°C, 5% CO_2_ in RPMI (Gibco) supplemented with 7.5% newborn calf serum (NCS; Sigma).

### Antibodies

The following antibodies have been used in this study: anti RhoA from Biomol No. NB-26007 and Abcam No. ab54835; anti-actin from Sigma No. A2228-100UL, anti-3x-Flag from Sigma No. F3165-1MG, anti-beta lactamase (TEM) from Abcam No. ab12251, anti-GFP from Sigma No. 11814460001, and anti-IgG from Cell Signalling No. 7076S.

### Cloning, expression and purification of recombinant CNF_Y_ variants

For crystallography purposes, truncated constructs were generated comprising a fragment lacking the catalytically active C-terminal domain (CNF_Y_1-704), a construct containing both C-terminal domains D4-5 (CNF_Y_526-1014) and another containing only the catalytic domain D5 (CNF_Y_720-1014). For crystallization of the full-length protein, a construct containing the inactive C866S variant of CNF_Y_ was produced.

The coding sequences of D4-5 (CNF_Y_1-704) and D5 (CNF_Y_720-1014) were both cloned into pET28c containing sequences coding for an N-terminal hexa-histidine tag and a thrombin protease cleavage site. The constructs were transformed into *E. coli* BL21 (DE3) (CNF_Y_1-704) or Rosetta II (DE3) (CNF_Y_720-1014). Native protein was expressed in lysogenic broth (LB) medium at 20°C after induction with 0.5 mM isopropyl-β-D-thiogalactopyranosid (IPTG) for 16-18 h (D1-4; CNF_Y_1-704) or 4 h (D5; CNF_Y_720-1024) (5), respectively. Seleno-L-methionine (Se-Met) labeled protein of CNF_Y_1-704 (D1-4) was expressed using M9 minimal medium.

After harvesting, the cell pellets were resuspended in lysis-buffer (for D1-4/CNF_Y_1-704: 1 x PBS, 400 mM NaCl, 5 mM β-mercaptoethanol, 5 mM MgSO_4_, 10 mM imidazole; for D5/CNF_Y_720-1014: 50 mM Tris/HCl pH 8.0, 400 mM NaCl, 5 mM imidazole) and lysed by sonification. The supernatant after centrifugation was mixed with 1 ml Ni-NTA resin pre-equilibrated with wash I buffer (D1-4/CNF_Y_1-704: 1 x PBS, 400 mM NaCl, 10 mM imidazole, 5 mM MgSO4, 5 mM β-mercaptoethanol; D5/CNF_Y_720-1014: 50 mM Tris/HCl pH 8, 400 mM NaCl, 5 mM imidazole) and incubated for 1 h on an overhead-shaker at 4°C. After washing with wash I buffer and wash II buffer (D1-4/CNF_Y_1-704: 1 x PBS, 400 mM NaCl, 20 mM imidazole, 5 mM MgSO_4_, 5 mM β-mercaptoethanol; D5/CNF_Y_720-1014: 50 mM Tris/HCl pH 8, 400 mM NaCl, 20 mM imidazole), elution of the protein was carried out with 12 x 1 ml of elution buffer (D1-4/CNF_Y_1-704: 1 x PBS, 400 mM NaCl, 250 mM imidazole, 5 mM MgSO4, 5 mM β-mercaptoethanol; D5/CNF_Y_720-1014: 50 mM Tris/HCl, 250 mM NaCl, 250 mM imidazole). Buffer exchange and tag cleavage with thrombin (1:50 mg/mg) were achieved over night by dialysis at 4°C in wash I buffer. To remove cleaved His-Tag, uncleaved protein and the thrombin protease, 1 ml of Ni-NTA resin and 5 ml of benzamidine-sepharose resin, respectively were mixed with the dialyzed protein solution. The collected flow-through predominantly contained pure protein. Further purification was achieved by size-exclusion chromatography. D1-4 (CNF_Y_1-704) was purified using a HiLoad 16/600 Superdex 200 pg (GE Healthcare) pre-equilibrated in buffer containing 20 mM Tris pH 8.0, 150 mM NaCl, 5 mM DTT. D5 (CNF_Y_720-1014) was purified using a HiLoad 16/600 Superdex 75 pg (GE Healthcare) pre-equilibrated in buffer containing 25 mM Tris pH 8.0, 100 mM NaCl. The proteins were then concentrated to 20 mg/ml, flash-frozen in liquid nitrogen and stored or directly used for crystallographic screens.

The gene encoding for the full-length protein of the CNF_Y_ C866S variant was cloned into a modified pCOLA Duet-1 vector (Novagen) encoding for an N-terminal Strep-tag II and TEV-protease recognition site (construct: CNF_Y_ C866S). In the case of D4-5 (CNF_Y_ 526-1014), the insert was amplified from pCNF_Y_3xFlag as template so that the three C-terminal FLAG-epitopes were included in the insert and cloned into the same modified pCOLA Duet-1 vector that was also used for the full-length toxin (construct: pVP-CNF_Y_526-1014-3xFlag). Both proteins were heterologously expressed in *E. coli* BL21 (DE3) in ZYM-5052 auto-inducing medium (Studier, 2005) at 20°C for 20-24 h.

In the case of D4-5 (CNF_Y_526-1014), the cell pellet was resuspended in a buffer containing 20 mM HEPES/NaOH pH 7.5, 300 mM NaCl, 2 mM TCEP, one tablet of complete EDTA-free protease inhibitor cocktail (Roche) and lysed by sonication. The protein was isolated from the supernatant after centrifugation for 1 h at 100.000 x g using a self-packed 10 ml column with Strep-Tactin Superflow High Capacity resin (IBA) and eluted from the column with a single step of 5 mM d-desthiobiotin. The affinity tag was cleaved off with TEV protease (1:50 mg/mg) at 4°C overnight. Gel filtration was carried out using a HiLoad 16/600 Superdex 200 pg column (GE Healthcare) in 20 mM HEPES/NaOH pH 7.5, 300 mM NaCl, 2 mM TCEP. The peak fractions were concentrated to 5 mg/ml and flash-frozen in liquid nitrogen for crystallization screening.

For the full-length protein, the cell pellet was resuspended in a buffer containing 20 mM HEPES/NaOH pH 7.5, 100 mM NaCl, 1 mM TCEP, one tablet of complete EDTA-free protease inhibitor cocktail (Roche) and lysed by sonication. The protein was isolated from the supernatant after centrifugation for 1 h at 100.000 x g using a self-packed 10 ml column with Strep-Tactin Superflow High Capacity resin (IBA) and eluted from the column with a single step of 5 mM d-desthiobiotin. The affinity tag was cleaved off with TEV protease (1:50 mg/mg) at 4°C overnight. Gel filtration was carried out using a HiLoad 16/600 Superdex 200 pg column (GE Healthcare) in 20 mM HEPES/NaOH pH 7.5, 100 mM NaCl, 1 mM TCEP. The fractions corresponding to the second peak in the chromatogram (elution volume 70-75 ml) were pooled and subjected to further size exclusion chromatography on a Superdex 200 Increase 10/300 GL column (GE Healthcare) in the same buffer. The peak fractions were concentrated to 27.5 mg/ml and flash-frozen in liquid nitrogen for crystallization screening. All chromatographic steps were carried out using an Äkta Purifier system (GE Healthcare). The samples were analyzed by SDS-PAGE (12%), and protein concentrations were determined from the absorbances at 280 nM with the extinction coefficients as calculated by Protparam (Gasteiger *et al*, 2003).

CNF_Y_-TEM fusion proteins were obtained in a similar manner after cloning into pET28a, overexpression in *<u>E. coli</u>* Rosetta II (DE3) and a two-step purification protocol involving Ni-NTA affinity and size exclusion chromatography.

### Crystallization

Crystallization trials were set up at room temperature with a HoneyBee 961 crystallization robot (Digilab Genomic Solutions) in Intelli 96-3 plates (Art Robbins Instruments) with 200 nl protein solution at different concentrations and 200 nl reservoir solution. Native D1-4 (CNF_Y_1-704) was crystallized in 0.1 M Tris pH 7.3-7.9, 0.2 M ammonium sulfate and 19-21% (w/v) PEG 5000 MME. The Se-Met derivative of D1-4 (CNF_Y_1-704) was crystallized in 0.1 M tri-sodium citrate pH 5.9-6.2, 0.2 M ammonium acetate and 28-32% (w/v) PEG 4000. Macro-seeding was applied in order to obtain well-diffracting crystals. As all tested compounds for cryoprotection were not tolerated by the samples, the crystals were flash-cooled without any additional cryoprotection. The catalytic domain D5 (CNF_Y_720-1014) yielded crystals in several PEG or ammonium sulfate containing conditions and the best diffracting crystals were obtained in 0.2 M ammonium fluoride with 20% (w/v) PEG 3350. Crystals were cryo-protected with either 25% glycerol or 100% Type A oil (Hampton Research) prior to flash freezing in liquid nitrogen. A single well-diffracting crystal of D4-5 (CNF_Y_526-1014) was obtained in the presence of 1 mM ATP in a condition containing 0.24 M magnesium chloride, 22.5% (w/v) PEG 2000 monomethyl ether. The crystal was harvested after 130 days of growth and cryo-protected by addition of 10% (v/v) (2*R*,3*R*)- 2,3-butanediol. A single crystal of sufficient diffraction quality of full-length CNF_Y_C866S was obtained in 1.4 M ammonium sulfate, 0.13 M lithium acetate, 0.1 M HEPES/NaOH pH 7.1. The crystal was harvested after 21 days of growth and after removal of satellite crystals cryo-protected by addition of 10% (v/v) (2*R*,3*R*)-2,3-butanediol.

### Data collection and processing

Data collection of native and Se-Met-derivatized D1-4 (CNF_Y_ 1-704) was performed on beamline PXIII of the Swiss Light Source (Paul Scherrer Institute, Villigen, Switzerland) and BESSY BL14.1 (Helmholtz Zentrum Berlin, Germany) (Mueller *et al*, 2015). High-resolution data of D5 (CNF_Y_ 720-1014) were recorded at beamline BL 14.2 of the BESSY II (Helmholtz-Zentrum Berlin, Germany). Datasets of domain D4-5 (CNF_Y_ 526-1014) and full-length CNF_Y_C866S were measured at beamline X06DA (PXIII) at the Swiss Light Source (Paul Scherrer Institute, Villigen, Switzerland). Data processing was achieved either manually via the XDS software package (Kabsch, 2010) or by using the AutoPROC (Vonrhein *et al*, 2011) toolbox (Global Phasing) executing XDS (Kabsch, 2010), Pointless (Evans, 2006), and Aimless (Evans & Murshudov, 2013). All datasets were recorded at a temperature of 100 K.

### Structure determination, refinement and model building

The structure of domain D1-3 (CNF_Y_ 1-704) was solved by single anomalous dispersion (SAD) using data collected at the selenium absorption edge. The initial phases were calculated using AutoSol (Terwilliger *et al*, 2009) and a partial model was generated running AutoBuild (Terwilliger *et al*, 2008), both components of the Phenix software package (Adams *et al*, 2010). The output model was analyzed in Coot (Emsley *et al*, 2010) and misplaced main chains were removed or corrected manually in order to obtain a reliable search-model for the following molecular replacement procedures against the dataset of native D1-4 (CNF_Y_ 1-704) and the full-length C866S variant. The structure of D5 (CNF_Y_ 720-1014) was determined by molecular replacement using the structure of the catalytic C-terminal domain of CNF1 from *E. coli* (PDB: 1HQ0, (Buetow *et al*, 2001)) as search-model. The structure of domain D4-5 (CNF_Y_526-1014) was determined by molecular replacement using the structure of domain D5 (CNF_Y_ 720-1014) and the region comprising residues 526-704 from domain D1-4 (CNF_Y_ 1-704). Phases for full-length CNF_Y_C866S were obtained by using both domain D1-4 (CNF_Y_1-704) and D5 (CNF_Y_720-1014) as search-models in molecular replacement. The molecular replacement procedures were carried out using Phaser (McCoy *et al*, 2007) from the Phenix suite (Adams *et al*, 2010). The structural models were built using Coot (Emsley *et al*, 2010) and crystallographic refinement was performed with Phenix.refine (Afonine *et al*, 2012) including the addition of hydrogens in riding positions and TLS-refinement. 5% of random reflections were flagged for the calculation of R_free_. The model of domain D1-4 (CNF_Y_ 1-704) was at 3.3 Å resolution and refined to R/R_free_ of 24/27% in space group P2_1_. The structure of domain D5 (CNF_Y_720-1014) was at 1.1 Å resolution and refined to R/R_free_ of 17/18% in space group P2_1_. The structure of domain D4-5 (CNF_Y_526-1014) was at 1.8 Å resolution and refined to R/R_free_ of 16/19% in space group P2_1_2_1_2_1_. The structural model of the full-length C866S variant of CNF_Y_ was at 2.7 Å resolution and refined to R/R_free_ of 21/24% in space group I2_1_2_1_2_1_. Data collection and refinement statistics are summarized in Tab EV1. Figures of crystal structures were prepared using the PyMOL Molecular Graphics System version 2.0.0 (Schrödinger, LLC).

### Construction of fusion plasmids and C-terminal *cnfY* deletions

To construct the plasmids for CNF_Y_ fusion proteins, the *blaM* gene and the 3xFlag tag were amplified using primers listed in Appendix Tab S4. pFU189 was used as a backbone from which the *luxCDABE* operon was removed by digestion with *Pst*I and *Not*I after which *blaM* or the 3xFlag tag were ligated into the vector, resulting in pTEM and p3xFLAG, respectively.

The PCR fragments of C-terminal *cnfY* deletions containing the *cnfY* promoter region were cloned into the *Bam*HI and *Pst*I sites of pTEM and p3xFLAG using the Quick-Fusion cloning kit (Biotool) with primers listed in Appendix Tab S4. For the construction of pCNF_Y_-GFP, *gfp* was excised from pFU31 and ligated into the *Pst*I and *Not*I sites of digested pCNF_Y_-TEM. All clones were transformed into *E. coli* DH10β and confirmed by sequencing. The plasmids were electroporated into *Y. pseudotuberculosis* YP147 (Δ*cnfY*) and selected for on LB agar plates containing the appropriate antibiotics.

### Site-directed mutagenesis of *cnfY*

Single residue mutants were generated by site-directed mutagenesis with primers listed in Appendix Tab S4. pCNF_Y_-TEM, pCNF_Y_-3xFLAG were used as templates. Clones were selected on LB containing the appropriate antibiotics. Mutations were verified by DNA sequencing.

### Detection of fusion proteins by immunoblotting

For protein expression, strain YP147 harboring the overexpression plasmids encoding full-length CNF_Y_ or the deletion variants was grown in BHI at 25°C overnight. Cells were harvested by centrifugation at 6,500 rpm and 4°C for 5 min. Cell pellets were washed with PBS and resuspended with lysis buffer (50 mM Tris-HCl pH 7.5, 100 mM NaCl, 5 mM MgCl_2_, 0.3% Triton X-100, 3 mg/ml lysozyme and protease inhibitor cocktail). After incubation at room temperature for 1 h, protein samples were centrifuged for 10 min and supernatants were sterilized with a 0.2 μm filter. To detect proteins, Western blot analysis was performed. The proteins were separated on a 10% SDS-polyacrylamide gel and transferred onto an Immobilon PVDF membrane (Millipore). Membranes were blocked in 5% BSA/TBST at 4°C overnight. Subsequently, the membrane was washed and incubated with primary antibody diluted in 5% BSA/TBST (1:10,000 anti-Flag (Sigma-Aldrich) or anti-Beta lactamase (Abcam)) at room temperature for 1 h. After washing, the secondary antibody diluted in 5% skim milk/TBST (1:5,000 anti-mouse IgG HRP (Cell Signaling Technology)) was added for 30 min at room temperature. After washing the membrane, proteins were visualized using the Western Lightning ECL II Kit (Perkin Elmer) and exposed on X-Ray film (GE Healthcare Amersham Hyperfilm ECL, Fisher Scientific).

### Nitrocefin secretion assay

Bacteria were grown overnight at 25°C in BHI containing the appropriate antibiotics. Subsequently, equal amounts of bacteria were pelleted by centrifugation at 13,000 rpm for 10 min. 95 μl of each supernatant was transferred to a 96-well plate in triplicate. 5 μl nitrocefin (2 mM) were added to each well and the plate was incubated at room temperature for 30 min. Beta-lactamase activity was determined at 390 nm (yellow) and 486 nm (red) using a VarioSkan plate reader (Thermo Scientific).

### Microbial viability assay

The microbial viability was assessed in equalized bacterial cultures using the BacTiter-Glo^TM^ Microbial Cell Viability Assay kit (Promega) according to the manufacturer’s recommendations and luminescence was measured using a VarioSkan plate reader (Thermo Scientific).

### Fluorescent actin staining

HEp-2 cells were seeded onto coverslips at a concentration of 5×10^4^ cells/well and allowed to attach overnight. The next day, cells were washed and incubated with an equal amount of cleared bacterial cell lysates for 24 h at 37°C, 5% CO_2_. After washing with PBS, cells were fixed in 4% paraformaldehyde for 15 min at room temperature. Subsequently, washed cells were permeabilized with 0.1% Triton X-100 in PBS for 1 min. The actin cytoskeleton was stained with FITC- or TRITC-Phalloidin (0.5 μg/ml in PBS; Sigma-Aldrich) and mounted on slides using ProLong® Gold Antifade mounting medium containing DAPI (Thermo scientific). Cells were visualized by fluorescence microscopy using an Axiovert II inverted fluorescence microscope (Carl Zeiss) with Axiocam HR and the AxioVision program (Carl Zeiss).

### CNF_Y_ translocation assay

In order to study the CNF_Y_ translocation into the host cells, a β-lactamase (TEM) reporter assay was performed using the LiveBLAzer-FRET B/G Loading Kit (Life Technologies). HEp-2 cells were seeded in 8-well μ-slides (Ibidi) at a concentration of 1.7×10^4^ cells/well and allowed to attach overnight. The next day, cells were washed and incubated for 24 h at 37°C, 5% CO_2_ with 20 µg/ml cleared lysates of *Y. pseudotuberculosis* expressing the different CNF_Y_-TEM fusion proteins. Cells were washed with PBS, followed by the addition of fresh media containing 20 mM HEPES. Cells were then stained with loading dye according to the manufactureŕs protocol. After staining for 1 h at room temperature, translocation was visualized by fluorescence microscopy using an Axiovert II with Axiocam HR and the AxioVision program (Carl Zeiss).

For the analysis of the translocation dynamics of CNF_Y_ 1-1014 and CNF_Y_ 1-526, surface-attached HEp-2 cells were charged with CCF4-AM in the dark for 1 h at 37°C and the change of fluorescence was immediately followed after addition of the respective CNF_Y_-TEM fusion construct in *Y. pseudotuberculosis* lysates (20 µg/ml) or the purified recombinant proteins (200 nM), using a CLARIOstar Plus fluorometer (BMG Labtech) or a Cytation 5 plate reader (Biotek), respectively.

### Fluorescence microscopy to visualize endocytosis

To test whether the CNF_Y_ deletion constructs are able to enter the cells through the endocytotic pathway, CellLight® Early or late Endosomes-RFP, BacMam 2.0 (Thermo Scientific) were used to investigate toxin entry to the host cells. HEp-2 cells were seeded 5×10^4^ cells/ml onto coverslips in 24 well plates and allowed to attach overnight. The next day, CellLight® reagent was added to HEp-2 cells around 20 particles per cells for 16 h and the cells were then incubated with CNF_Y_ toxin on ice for 30 min. Subsequently, cells were washed and transferred to 37°C for 30, 90 or 180 min. To investigate colocalization of CNF_Y_ with early or late endosome, cells were then fixed and visualized by using a fluorescence microscope (Axiovert II with Axiocam HR, Carl Zeiss) and the AxioVision program (Carl Zeiss).

### Biochemical analysis of RhoA deamidation

Cells were seeded in 10 cm cell culture dishes at the concentration of 2.2 x 10^6^ cells/dish and allowed to attach overnight. The next day, cells were washed and incubated for 4 h with 20 µg/ml of cleared lysates from *Y. pseudotuberculosis* expressing different CNF_Y_ derivatives at 37°C, 5% CO_2_. Cells were washed with cold PBS and lysed in 150 μl lysis buffer containing 50 mM Tris-HCl (pH7.4), 100 mM NaCl, 2 mM MgCl_2_, 10% NP-40 and 0.5 mM phenyl-methyl-sulfonyl fluoride (PMSF). Cells were then scraped off and centrifuged for 30 min (13,000 rpm, 4°C). Sodium dodecyl sulfate (SDS) sample buffer was added to the clear lysates and samples were separated on 12% SDS-gel. After blotting onto a PVDF membrane, RhoA was detected using mouse anti-RhoA IgG (Millipore) (1:1000) as a primary antibody and followed by secondary antibody goat anti-mouse IgG-HRP (Cell signaling). Membranes were visualized using the Western Lightning ECL II Kit (Perkin Elmer) and exposed on X-Ray film.

### *In vitro* RhoA shift assay

*In vitro* RhoA shift assays were performed in order to check for proper folding and catalytic activity of CNF_Y_ deletion constructs. Cells were seeded on 150 mm dish at the concentration of 5 x 10^6^ cells/dish and allowed to attach overnight. The next day, cells were washed with PBS and lysed in 300 μl lysis buffer (50 mM Tris-HCl, pH 7.5, 5 mM MgCl_2_, 1 mM EDTA, 10% NP-40 and 1 mM dithiothreitol (DTT)). Cells were then scraped off and centrifuged 13,000 rpm at 4°C for 30 min. 20 µg/ml of cleared lysates from *Y. pseudotuberculosis* expressing different CNF_Y_ derivatives were added to the prepared cell extracts and then incubated for 4 h, or 1 µM purified CNF_Y_ or CNF_Y_ 526-1014 protein were incubated in the cell extracts for 10-60 min at 37°. The reactions were stopped by adding SDS-sample buffer and heated at 95°C for 10 min. Samples were then subjected to 12% SDS-PAGE. After blotting onto a PVDF membrane, blots were developed as mentioned above.

### Quantification of RhoA deamidation by mass spectrometry

Recombinant RhoA (20 µM) was incubated at 37°C with full-length CNF_Y_ or CNF_Y_ 526-1014 in 100 µl assay-buffer (20 mM HEPES/NaOH, 300 mM NaCl, 2 mM MgCl_2_, 2 mM TCEP, 100 µM GDP). Samples, after setup for 30 min, 3 h and 16 h, were immediately mixed with 2-fold SDS-PAGE loading buffer, denatured for 5 min at 95°C and subjected to SDS-PAGE by applying 2 µg of RhoA per lane. After staining with Coomassie Blue, bands corresponding to RhoA were cut out, destained with 30% acetonitrile in 50 mM TEAB and then dehydrated with 100% acetonitrile. After reduction with 20 mM TCEP in 50 mM TEAB (1 h, 56°C), alkylation was performed by 20 mM MMTS in 50 mM TEAB (1 h, RT), followed by in-gel digestion over night with 50 ng trypsin (Promega) in 50 mM TEAB. Peptides were extracted by using 10 gel volumes of 0.5% formic acid in 30% acetonitrile, vacuum dried and resuspended in 0.1% formic acid before applying to Evotips as recommended by the manufacturer (Evosep). Peptide sequencing of RhoA was carried out by tandem mass spectrometry on an Evosep One HPLC system linked with a timsTOF Pro mass spectrometer (Bruker). Separation of modified peptide ions was supported by ion mobility and RhoA deamidation in the peptide QVELALWDTAG<u>Q</u>EDYDR was identified by using PEAKS 10+ software. Deamidated and corresponding non-deamidated RhoA peptide variants were manually validated at the level of their distinct HPLC-retention times and corresponding representative fragmentation spectra (MS2; Data Analysis, Bruker). RhoA modification was then quantified by selected ion chromatograms (MS1; Data Analysis, Bruker). CNF_Y_-dependent deamidation was determined in comparison to non-treated RhoA samples (negative control) and based on the area under peaks for double charged non-deamidated peptides (m/z 1004.9645 ± 0.005) and deamidated peptides (retention time + 0.3 min, m/z 1005.4590 ± 0.005).

## Acknowledgements

We thank the beamline staff at the Helmholtz Centre Berlin (Germany) and the Paul Scherrer Institute (Villigen, Switzerland) for providing access to beamlines BL14.1 and BL14.2 at the BESSYII electron storage ring and to beamline X06DA at the SLS synchrotron. Experiments at the SLS have received funding from the European Union’s Horizon 2020 research and innovation program under grant agreement n.° 730872, project CALIPSOplus. Andrea Berger and Ute Widow are acknowledged for excellent technical assistance. We thank Janina Schweer for the construction of initial plasmids and strains. Manfred Nimtz and Josef Wissing helped with mass spectrometry experiments. PD and PC received funding from the SFB1009 (Project no. 194468054) of the German Research Foundation (DFG). TH and PC were supported by the HZI Graduate School for Infection Research.

## Author contributions

PD and WB conceived the project. PL, EMG and TH produced recombinant proteins and performed crystallization and structure determination. CR, PC, SM, and TS conducted work with *Y. pseudotuberculosis* and analyzed microscopy experiments with eukaryotic cells. Similar microscopy experiments with recombinant proteins were performed by AS, MK and SSD. WJB and LJ performed and analyzed mass-spectrometric experiments. PD, PL, and WB wrote the manuscript.

## Data availability

Coordinates and structure factor amplitudes have been deposited in the Protein Data Bank with accession codes 6Q7Z (CNF_Y_ 1-704), 6Q7Y (CNF_Y_ 720-1014), 6Q80 (CNF_Y_ 526-1014) and 6Q7X (CNF_Y_ 1-1014 C866S).

## Expanded View Tables

**Table EV1.**
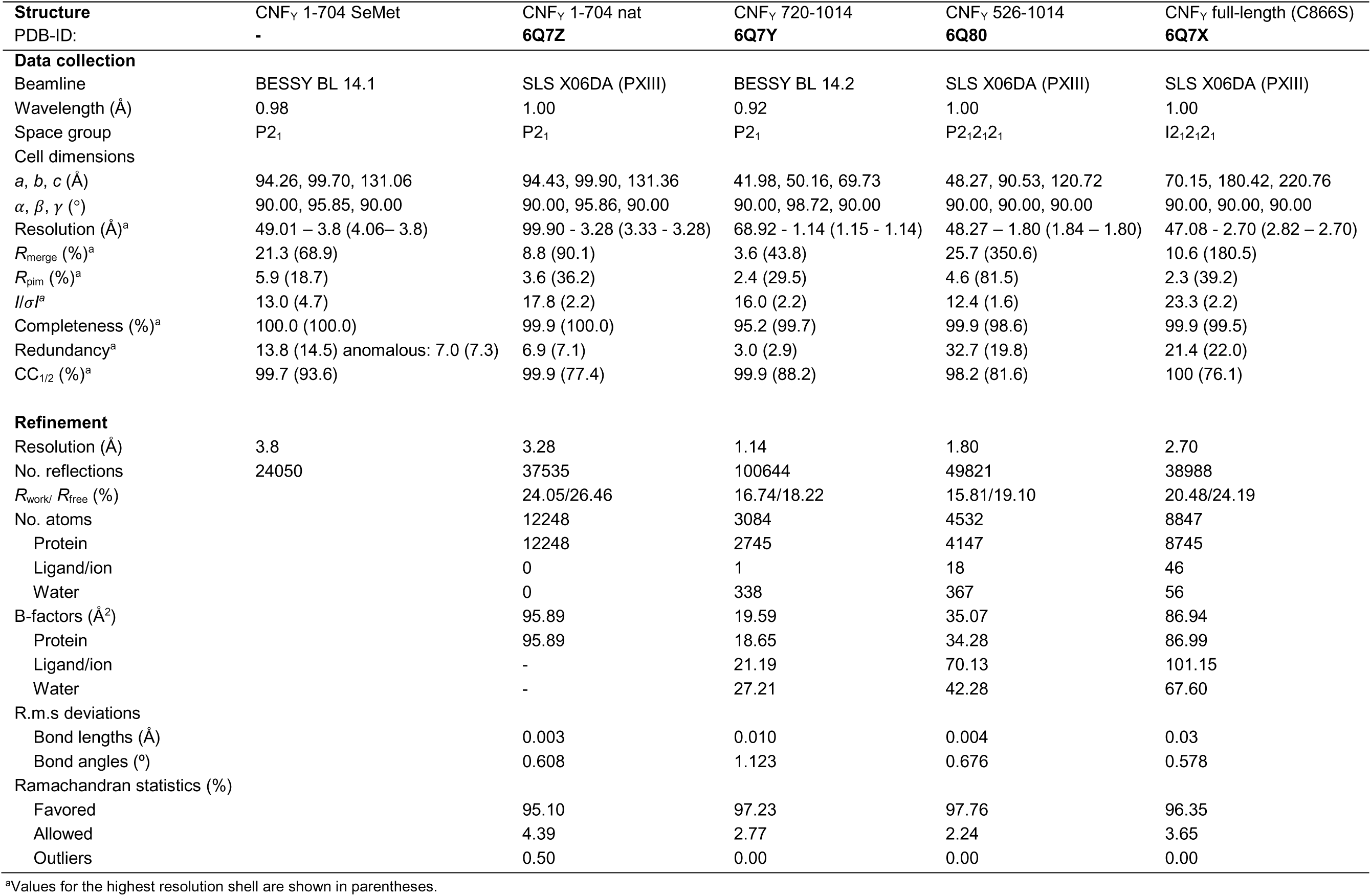
**X-ray data collection and refinement statistics.**

## Expanded View Figures

**Figure EV1.**
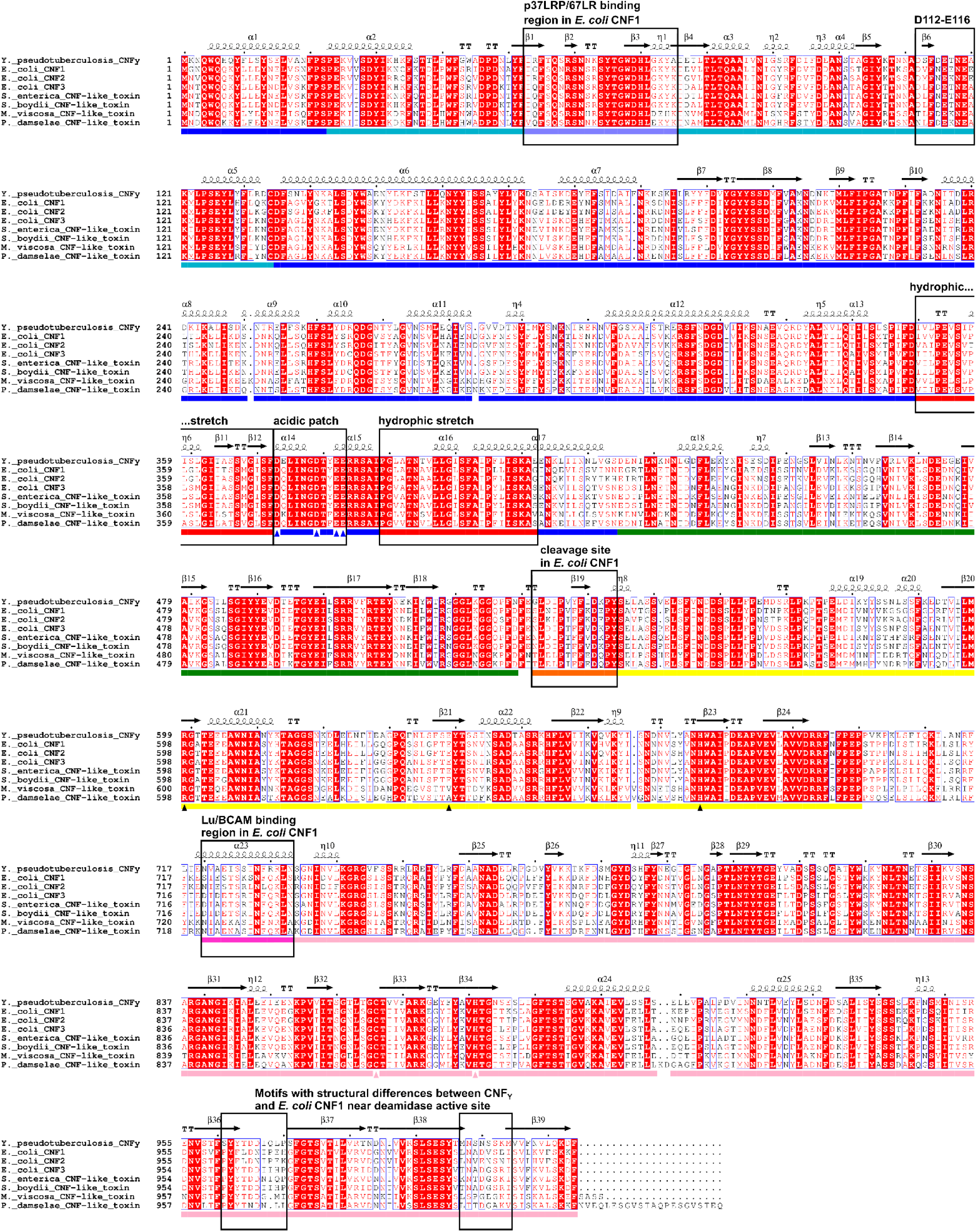
**Sequence alignment of CNF_Y_ to other CNFs.** Sequences from top to bottom: 1. CNF_Y_ from *Yersinia pseudotuberculosis*, 2. CNF1 from *E. coli* (NCBI accession: AAA85196, 61% sequence identity to CNF_Y_), 3. CNF2 from *E. coli* (NCBI accession: ACT33566, 61% sequence identity), 4. CNF3 from *E. coli* (NCBI accession: CAK19001, 68% sequence identity), 5. CNF from *Salmonella enterica* (NCBI accession: WP_079946821, 69% sequence identity), 6. CNF from *Shigella boydii* (NCBI accession: WP_075330563, 68% sequence identity), 7. CNF from *Moritella viscosa* (NCBI accession: AHI58923, 58% sequence identity). 8. CNF from *Photobacterium damselae* (NCBI accession: WP_005306733, 58% sequence identity). Columns with identical residues are highlighted in red. The sequence alignment was generated using ClustalX (Larkin, Blackshields et al., 2007) and formatted using the ESPript 3 webservice (Robert & Gouet, 2014). The alignment has been annotated with the secondary structure extracted from the structural model of full-length CNF_Y_. The domains of CNF_Y_ are indicated below the sequences using the coloring scheme used in Fig. 1. Specific residues that are supposed to be functionally relevant are marked by arrows in the domain annotation.

**Figure EV2.**
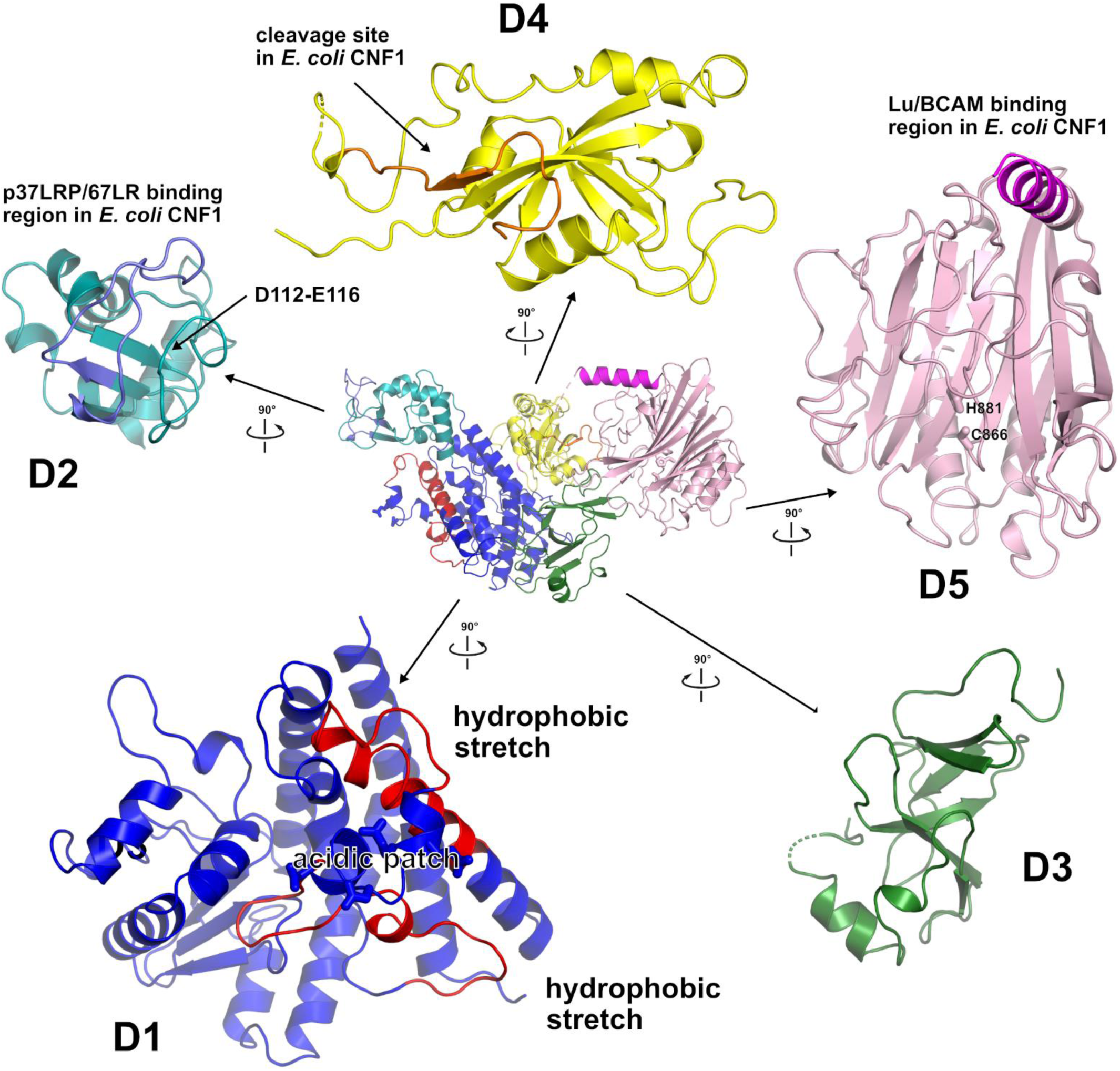
**Exploded view of CNF_Y_.** Cartoon representation of full-length CNF_Y_ (middle) surrounded by enlarged perpendicular views of the individual domains D1 to D5, colored according to domain boundaries determined with PiSQRD (Aleksiev *et al*, 2009). Dark blue: domain D1, cyan: domain D2, dark green: domain D3, yellow: ADP-ribosyltransferase-like domain D4, pink: deamidase domain D5. Other colors indicate the position of sequence motifs that have been identified in *E. coli* CNF1, namely light purple: p37LRP/67LR receptor-binding motif, red: hydrophobic stretches predicted to form membrane-inserting α-helices, orange: cleavage site, magenta: main Lu/BCAM receptor-binding motif.

**Figure EV3.**
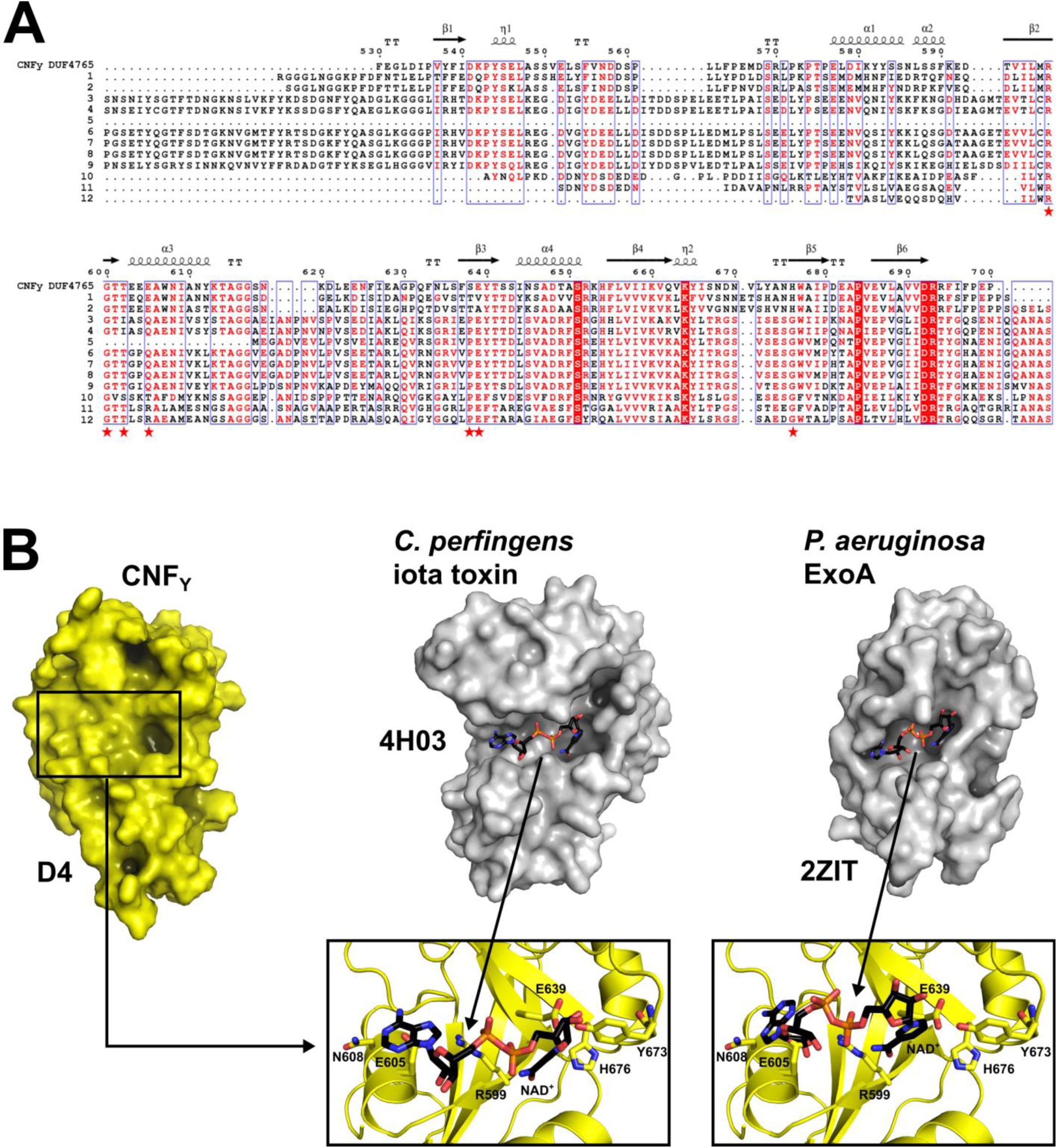
**Exploded view of CNF_Y_.** Cartoon representation of full-length CNF_Y_ (middle) surrounded by enlarged perpendicular views of the individual domains D1 to D5, colored according to domain boundaries determined with PiSQRD (Aleksiev *et al*, 2009). Dark blue: domain D1, cyan: domain D2, dark green: domain D3, yellow: ADP-ribosyltransferase-like domain D4, pink: deamidase domain D5. Other colors indicate the position of sequence motifs that have been identified in *E. coli* CNF1, namely light purple: p37LRP/67LR receptor-binding motif, red: hydrophobic stretches predicted to form membrane-inserting α-helices, orange: cleavage site, magenta: main Lu/BCAM receptor-binding motif.

**Figure EV4.**
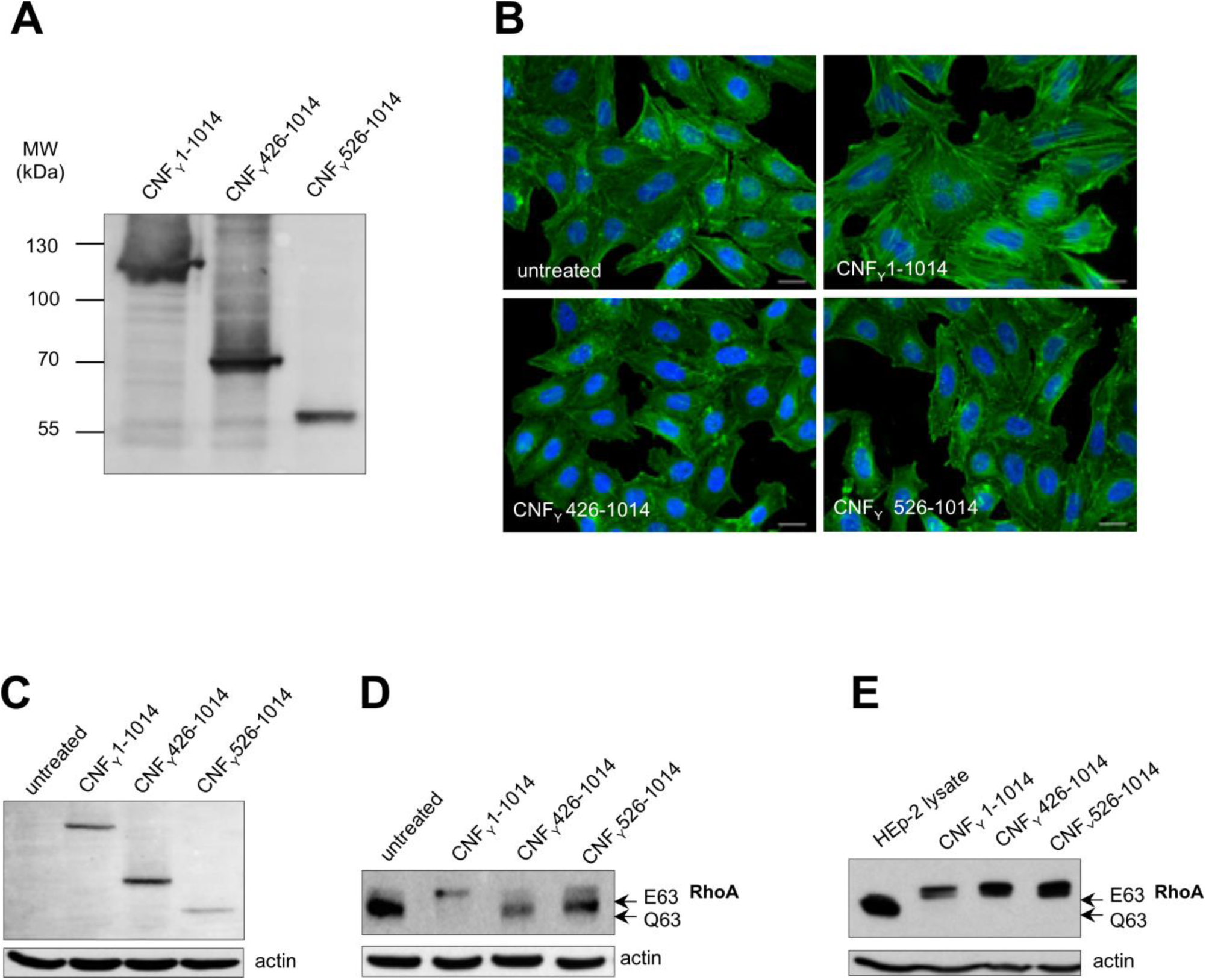
**C-terminal domain D3-5 and D4-5 are able to bind to host cells and deaminate RhoA.** A The expression of purified recombinant CNF_Y_ N-terminal domains D3-5 (CNF_Y_ 426-1014) or D4-5 (CNF_Y_ 526- 1014). B HEp-2 cells remain untreated or were incubated with 500 nM purified full-length CNF_Y_, domains D3-5 (CNF_Y_ 426-1014) or D4-5 (CNF_Y_ 526-1014) for 24 h. The formation of large, multinuclear cells was observed by fluorescence microscopy. The cell nuclei were strained with DAPI (blue) and the actin cytoskeleton was stained using FITC-Phalloidin (green). The white scale bar is 20 µm. C Binding of purified full-length CNF_Y_, domains D3-5 (CNF_Y_ 426-1014) or D4-5 (CNF_Y_ 526-1014) to HEp-2 cells was analyzed by immunoblotting after HEp-2 cells were incubated with 500 nM of the purified toxin or toxin fragments for 4 h. D HEp-2 cells treated with 500 nM purified full-length CNF_Y_, protein fragments D3-5 (CNF_Y_ 426-1014) or D4-5 (CNF_Y_ 526-1014) were lysed and the deamidation of RhoA was analyzed by the mobility shift of the modified GTPase detected by immunoblotting. E The activity of purified CNF_Y_ and the protein fragments D3-5 (CNF_Y_ 426-1014) or D4-5 (CNF_Y_ 526-1014) was tested by analyzing the deamidation of RhoA in HEp-2 cell lysates by the mobility shift of the modified GTPase detected by immunoblotting.

## Appendix Tables

**Table S1.**
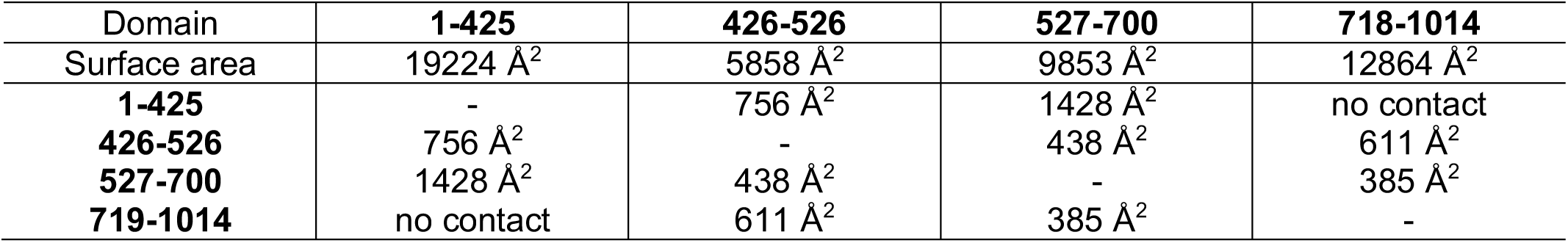
**Domain Interfaces.** The CNF_Y_ structural model was split at the domain boundaries and surfaces of the isolated domains and domain-domain interface areas were calculated using PDBe PISA (Krissinel & Henrick, 2007).

| Domain | **1-425** | **426-526** | **527-700** | **718-1014** |
| --- | --- | --- | --- | --- |
| Surface area | 19224 Å<sup>2</sup> | 5858 Å<sup>2</sup> | 9853 Å<sup>2</sup> | 12864 Å<sup>2</sup> |
| **1-425** | - | 756 Å<sup>2</sup> | 1428 Å<sup>2</sup> | no contact |
| **426-526** | 756 Å<sup>2</sup> | - | 438 Å<sup>2</sup> | 611 Å<sup>2</sup> |
| **527-700** | 1428 Å<sup>2</sup> | 438 Å<sup>2</sup> | - | 385 Å<sup>2</sup> |
| **719-1014** | no contact | 611 Å<sup>2</sup> | 385 Å<sup>2</sup> | - |

**Table S2.** **Structural homology of domain D4.** Structures with folds similar to domain D4 of CNF_Y_ as identified by DALI (Holm & Rosenstrom, 2010). Listed below are the 10 highest scoring results, ordered according to their Z-score in descending order.

| PDB entry | Chain | Z-score | C<sub>α</sub><br>R.m.s.d.<br>(Å) | length of<br>alignment | number of<br>residues in<br>target | %<br>sequence-<br>identity | Description |
| --- | --- | --- | --- | --- | --- | --- | --- |
| 5LYH | A | 5.6 | 3.6 | 101 | 185 | 11 | Human poly(ADP-ribose) polymerase 14 |
| 1WFX | A | 5.5 | 2.9 | 78 | 180 | 19 | 2'-phosphotransferase from *Aeropyrum pernix* |
| 3HKV | A | 5.3 | 4.0 | 102 | 192 | 9 | Human poly(ADP-ribose) polymerase 10 |
| 4F0D | A | 5.2 | 3.8 | 103 | 238 | 12 | Human poly(ADP-ribose) polymerase 16 |
| 2Q5T | A | 5.2 | 4.1 | 100 | 605 | 11 | Cholix toxin from *Vibrio cholerae* |
| 2AUA | B | 4.7 | 3.0 | 72 | 210 | 13 | Protein of unknown function from *Bacillus cereus* |
| 1DDT | A | 4.7 | 3.6 | 94 | 523 | 12 | Diphtheria toxin from *Corynebacterium diphtheriae* |
| 4HYF | A | 4.5 | 4.7 | 101 | 227 | 10 | Human tankyrase-2 |
| 4TLV | A | 4.4 | 4.9 | 89 | 582 | 11 | CARDS toxin from *Mycoplasma pneumoniae* |
| 3AQ2 | A | 4.4 | 3.1 | 96 | 191 | 10 | Protein 6b from *Agrobacterium vitis* |

**Table S3:** Strains and plasmids.

| Strain | Characteristics | Source/Reference |
| --- | --- | --- |
| YPIII | Wild type *Yersinia pseudotuberculosis* serogroup III | (Bolin, Norlander et al., 1982) |
| YP147 | YPIII, $\Delta cnfY::Kn^R$ | (Schweer, Kulkarni et al., 2013) |
| DH10β | | Invitrogen |
| Rosetta II (DE3) |  | Novagen |
| BL21 (DE3) |  | Novagen |
| **Plasmid** |  |  |
| pFU189 | Cm<sup>R</sup> | (Uliczka & Dersch, 2012) |
| pTEM | Cm<sup>R</sup> | this study |
| pCNF<sub>Y</sub>-TEM | Cm<sup>R</sup> | this study |
| pCNF<sub>YC866S</sub>-TEM | Cm<sup>R</sup> | this study |
| pCNF<sub>Y1-719</sub>-TEM | Cm<sup>R</sup> | this study |
| pCNF<sub>Y1-526</sub>-TEM | Cm<sup>R</sup> | this study |
| pCNF<sub>Y1-443</sub>-TEM | Cm<sup>R</sup> | this study |
| pCNF<sub>YΔ39-134</sub>-TEM | Cm<sup>R</sup> | this study |
| pCNF<sub>YΔ134-426</sub>-TEM | Cm<sup>R</sup> | this study |
| pCNF<sub>YΔ39-426</sub>-TEM | Cm<sup>R</sup> | this study |
| pCNF<sub>YΔ527-719</sub>-TEM | Cm<sup>R</sup> | this study |
| pCNF<sub>YΔ527-699</sub>-TEM | Cm<sup>R</sup> | this study |
| pCNF<sub>YE382/383K</sub>-TEM | Cm<sup>R</sup> | this study |
| p3xFLAG | Cm<sup>R</sup> | this study |
| pCNF<sub>Y</sub>-3xFLAG | Cm<sup>R</sup> | this study |
| pCNF<sub>YC866S</sub>-3xFLAG | Cm<sup>R</sup> | this study |
| pCNF<sub>Y1-719</sub>-3xFLAG | Cm<sup>R</sup> | this study |
| pCNF<sub>Y1-526</sub>-3xFLAG | Cm<sup>R</sup> | this study |
| pCNF<sub>Y1-443</sub>-3xFLAG | Cm<sup>R</sup> | this study |
| pCNF<sub>YΔ39-134</sub>-3xFLAG | Cm<sup>R</sup> | this study |
| pCNF<sub>YΔ134-426</sub>-3xFLAG | Cm<sup>R</sup> | this study |
| pCNF<sub>YΔ39-426</sub>-3xFLAG | Cm<sup>R</sup> | this study |
| pCNF<sub>YΔ527-719</sub>-3xFLAG | Cm<sup>R</sup> | this study |
| pCNF<sub>YΔ527-699</sub>-3xFLAG | Cm<sup>R</sup> | this study |
| pCNF<sub>YE382/383K</sub>-3xFLAG | Cm<sup>R</sup> | this study |
| pCNF<sub>Y I535L/P536A/V537G</sub>-TEM | Cm<sup>R</sup> | this study |
| pCNF<sub>YI535L/P536A/V537G/F539L/D541A/K542A</sub>-TEM | Cm<sup>R</sup> | this study |
| pCNF<sub>Y</sub>-GFP | Cm<sup>R</sup> | this study |
| pCNF<sub>Y1-719</sub>-GFP | Cm<sup>R</sup> | this study |
| pCNF<sub>Y1-526</sub>-GFP | Cm<sup>R</sup> | this study |
| pCNF<sub>Y1-443</sub>-GFP | Cm<sup>R</sup> | this study |
| pCOLA-Duet-1 | Kan<sup>R</sup> | Novagen |
| pET28c-CNF<sub>Y1-704</sub> | Kan<sup>R</sup> - construct for crystallography purposes | this study |
| pET28c-CNF<sub>Y720-1014</sub> | Kan<sup>R</sup> - construct for crystallography purposes | this study |
| pVP-CNF<sub>Y526-1014</sub>-3xFlag | Kan<sup>R</sup> - construct for crystallography purposes | this study |
| pVP-CNF<sub>YFL-C866S</sub> | Kan<sup>R</sup> - construct for crystallography purposes | this study |

**Table S4:**
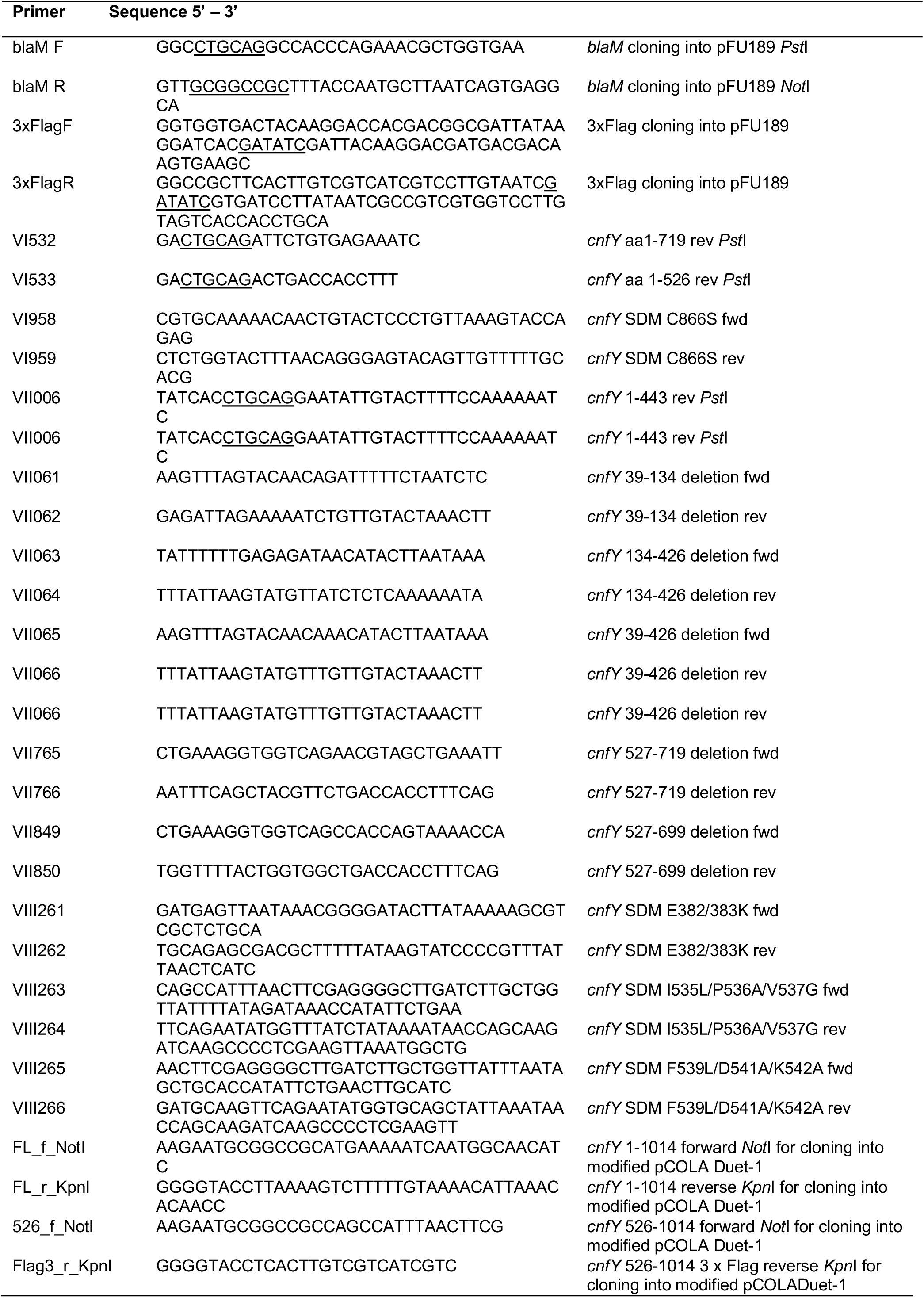
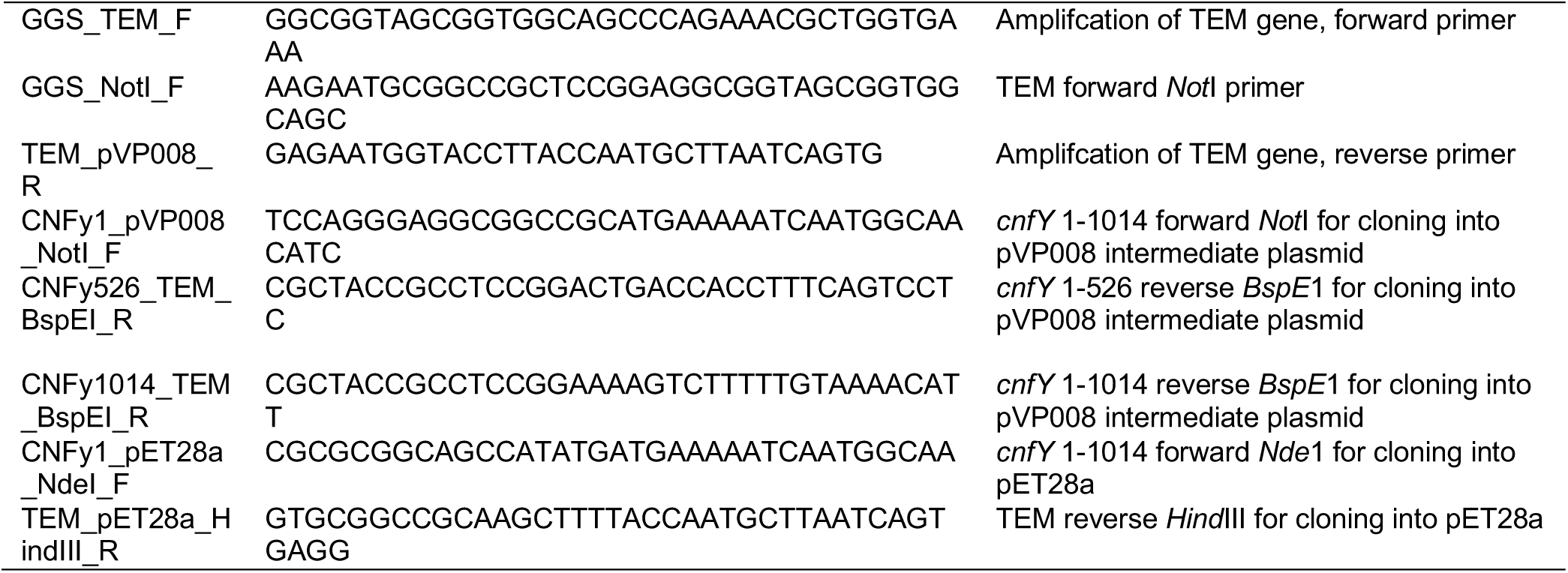
Oligonucleotide primers.

## Appendix Figures

**Figure S1.**
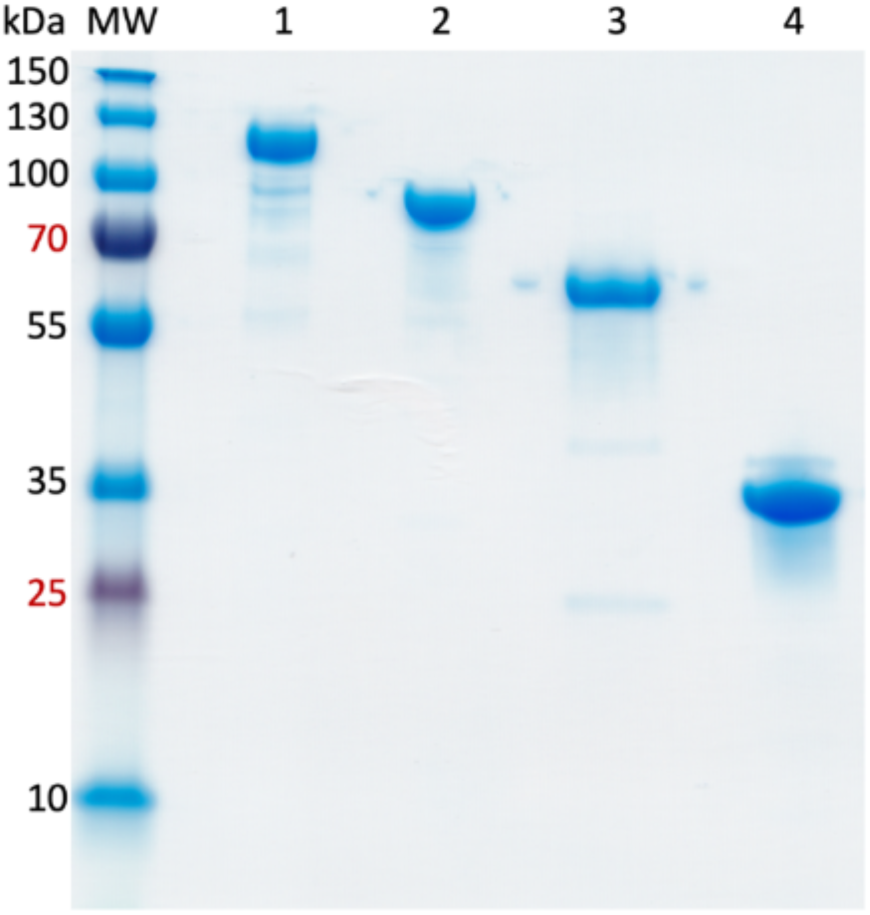
**SDS-PAGE of purified CNF_Y_ fragments for crystallization** Coomassie-stained SDS-PAGE (Any kD Mini-PROTEAN TGX, Bio-Rad). The fragments have been produced in *E. coli* and were purified as described in the methods section. MW: Molecular weight standard (PageRuler Plus prestained ladder, Thermo), 1: CNF**_Y_** full-length, 2: CNF**_Y_** 1-704, 3: CNF**_Y_** 526-1014, 4: CNF**_Y_** 720-1014. 2 µg of each construct have been loaded onto the gel.

**Figure S2.**
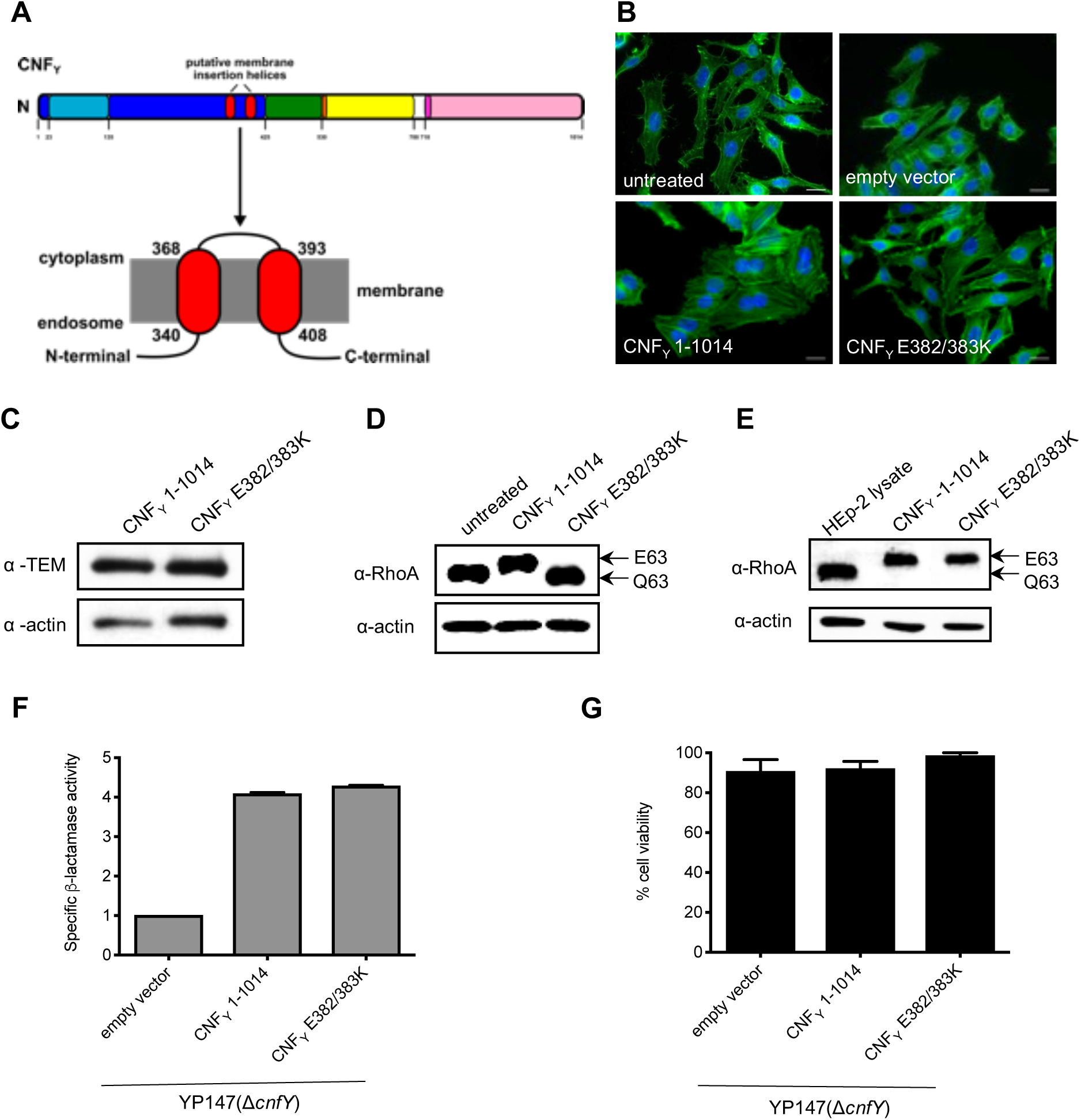
**Characterization of secretion, cell binding, and activity of the CNF_Y_ E382/383K variant. A** A schematic overview of CNF_Y_ illustrating the predicted helical regions of the D2 domain exposing the acidic loop. B HEp-2 cells were incubated with 20 µg/ml of whole cell extract of *Y. pseudotuberculosis* expressing CNF_Y_, the toxin variant CNF_Y_ E382/383K or no CNF_Y_ protein (harboring empty vector) for 24 h. The formation of large, multinuclear cells was observed by fluorescence microscopy. The cell nuclei were strained with DAPI (blue) and the actin cytoskeleton was stained using FITC-Phalloidin (green). The white scale bar is 20 µm. C The binding of CNF_Y_ to HEp-2 cells was analyzed by immunoblotting after HEp-2 cells were incubated with 20 µg/ml of whole cell extract of *Y. pseudotuberculosis* expressing CNF_Y_ for 4 h. D Cells incubated with 20 µg/ml of whole cell extract of *Y. pseudotuberculosis* expressing CNF_Y_ and the toxin variant CNF_Y_ E382/383K for 4 h were lysed and the deamidation of RhoA was analyzed by the mobility shift of the modified GTPase detected by immunoblotting. E The activity of the purified CNF_Y_ derivatives was tested by analyzing the deamidation of RhoA in HEp-2 cell lysates by the mobility shift of the modified GTPase detected by immunoblotting. F CNF_Y_-TEM and CNF_Y_ E382/383-TEM secretion was determined by measuring changes in absorbance of nitrocefin at 390 nm (yellow) and 486 nm (red). G The microbial viability of the bacteria expressing CNF_Y_ and CNF_Y_ E382/383 was assessed in equalized bacterial cultures using the BacTiter-Glo^TM^ Microbial Cell Viability Assay kit.

**Figure S3.**
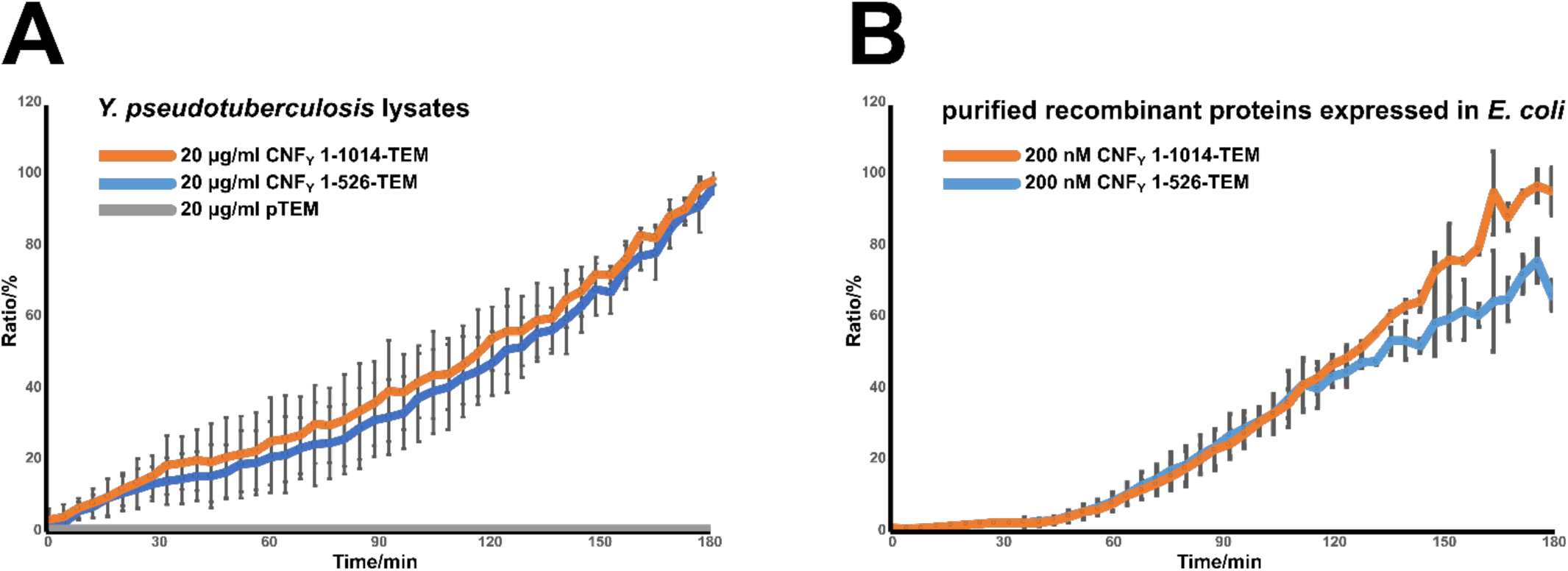
**Translocation efficiency of full-length CNF_Y_ vs. CNF_Y_ 1-526** HEp-2 cells were loaded with CCP4-AM and then incubated with the indicated CNF_Y_-TEM fusion proteins applied either as (A) lysates from *Y. pseudotuberculosis* or as (B) purified recombinant proteins purified from *E. coli*. The development of blue fluorescence, indicating translocation of TEM-β-lactamase into the cytosol, is plotted for 3 h, taking the final reading as 100%. Error bars represent standard deviations of (A) eight or (B) three replicates. Lysate from *Y. pseudotuberculosis* harboring just the pTEM plasmid served as a negative control and demonstrates that TEM-β-lactamase cannot enter HEp-2 cells when it is not fused to the translocation machinery (A).

**Figure S4.**
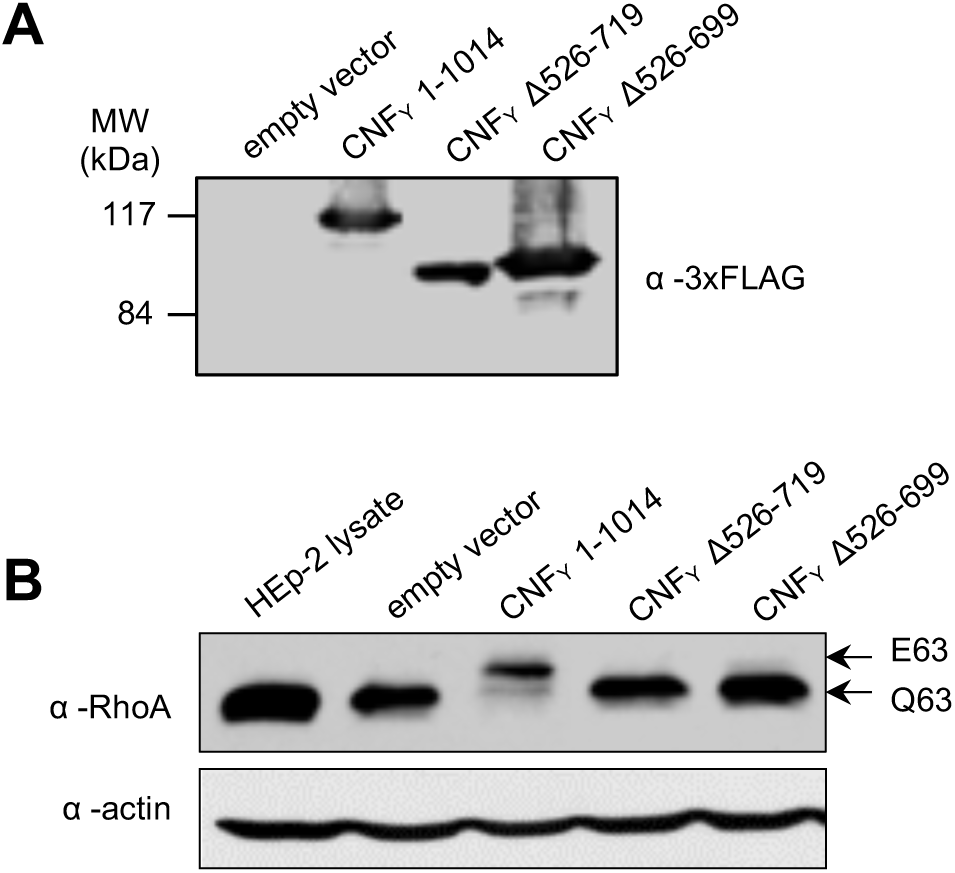
**Expression and analysis of CNF_Y_ Δ527-719 or CNF_Y_ Δ527-699 deletion proteins missing domain D3.** A Expression of 3xFlag-lagged version of CNF_Y_ wildtype protein (CNF_Y_1-1014 aa) and the internal deletion derivatives CNF_Y_ Δ526-699 and CNF_Y_ Δ526-719. B HEp-2 cell lysates were incubated with 20 µg/ml of whole cell extract of *Y. pseudotuberculosis* expressing the indicated CNF_Y_ protein and their activity was tested by analyzing the deamidation of RhoA in HEp-2 cell lysates by the mobility shift of the modified GTPase detected by immunoblotting.

**Figure S5.**
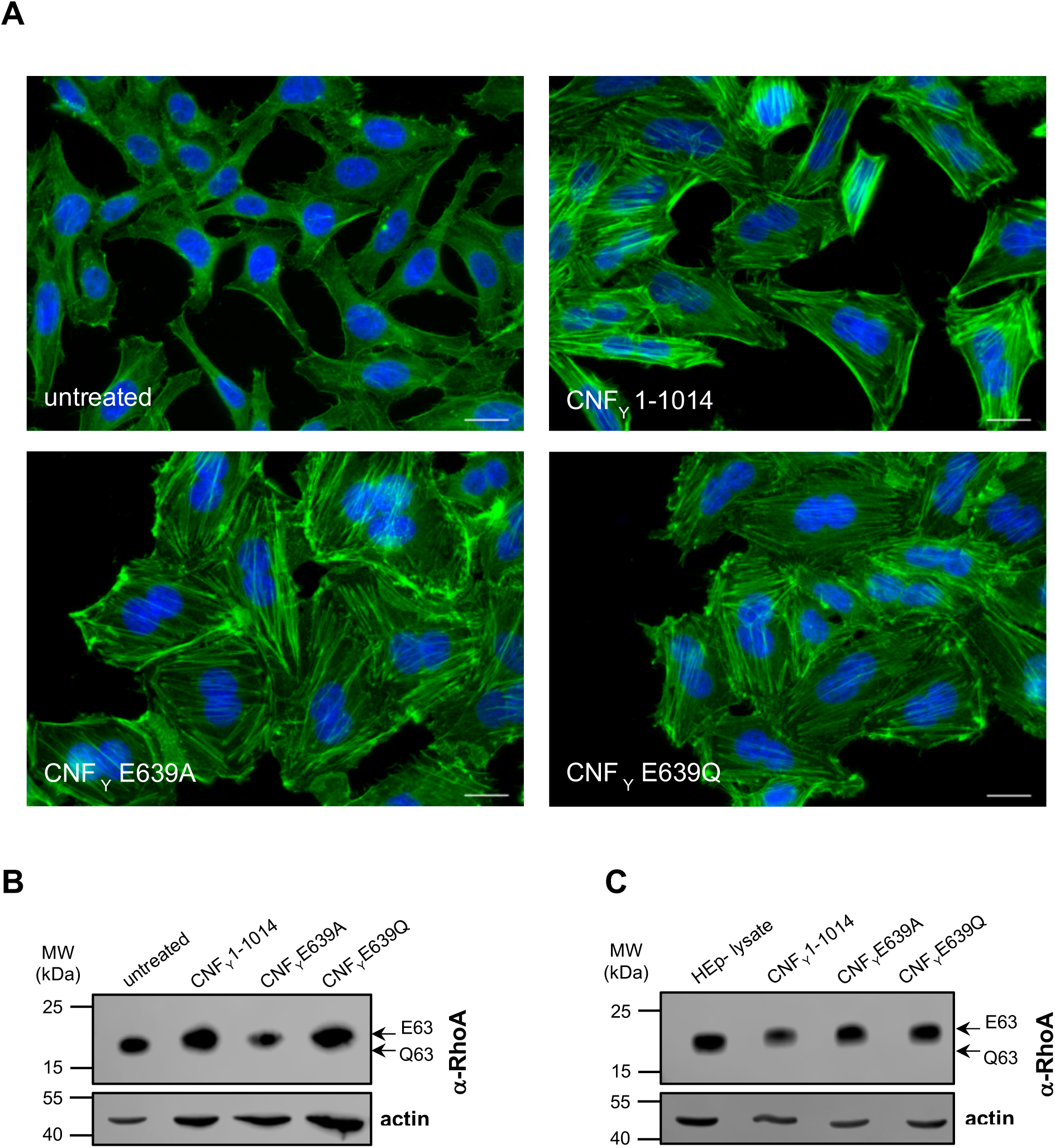
**Expression of CNF_Y_ E639A and 639Q and *in vitro* deamidation of RhoA.** A Purified CNF_Y_ E639A and 639Q proteins (500 nM) were added to HEp-2 cells for 24 h. Cell nuclei were strained with DAPI (blue) and the actin cytoskeleton was stained using FITC-phalloidin (green) after treatment of cells with CNF_Y_ toxins and cells assessed by microscopy. The white scale bar is 20 µm. B HEp-2 cells treated with 500 nM purified full-length CNF_Y_, CNF_Y_ CNF_Y_ E639A and 639Q were lysed and the deamidation of RhoA was analyzed by the mobility shift of the modified GTPase detected by immunoblotting. C The activity of the CNF_Y_ mutant proteins were tested by analyzing the deamidation of RhoA in HEp-2 cell lysates by the mobility shift of the modified GTPase detected by immunoblotting.

**Fig. S6:**
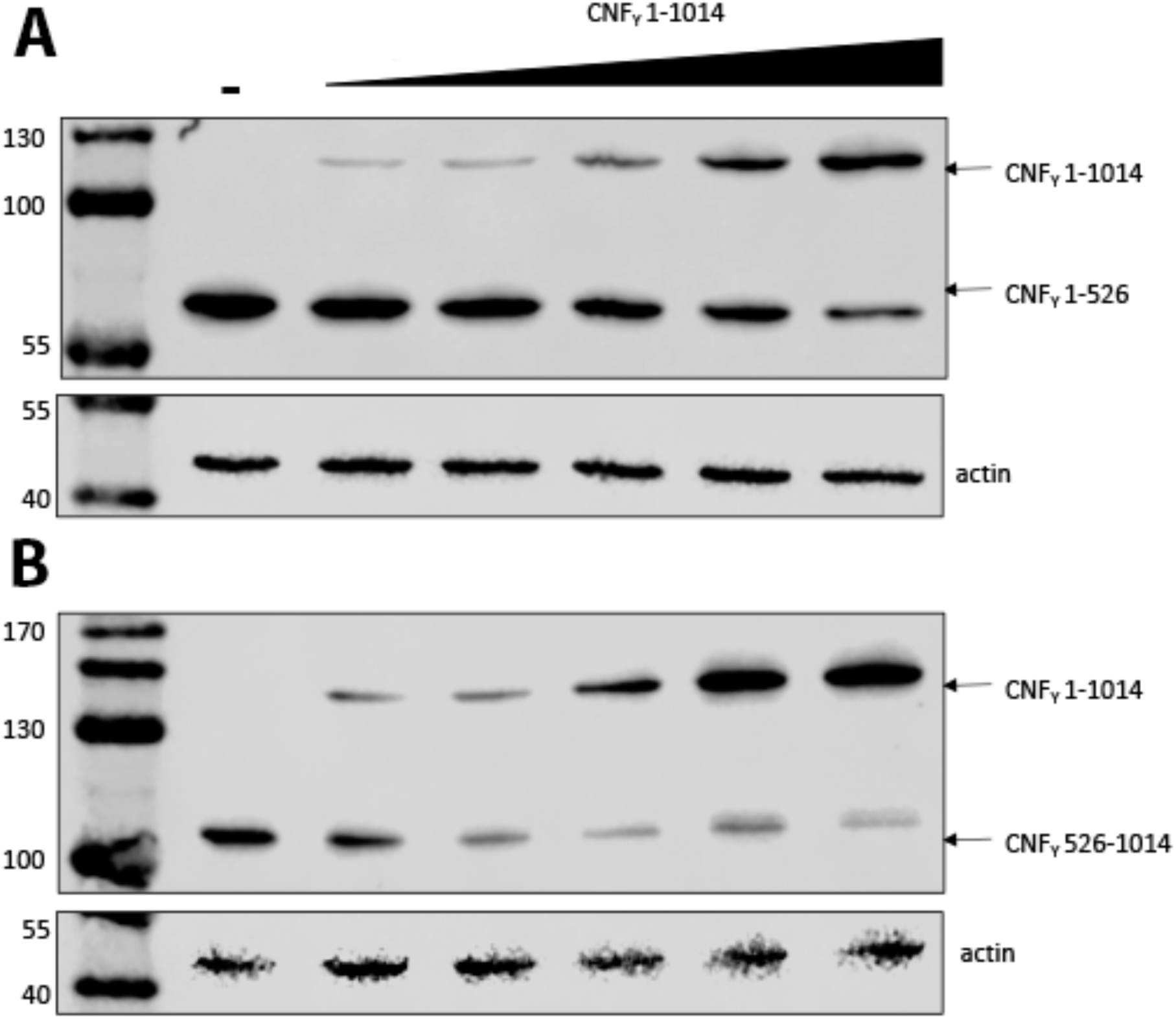
**Binding of the CNF_Y_ D1-3 and D4-5 domain can be efficiently replaced by full-length CNF_Y_.** HEp-2 cells were incubated with 25 µg/ml of 3xFlag-tagged CNF_Y_ 1-526 (**A**) or CNF_Y_ 526-1014 (**B**) and no (-) or increasing amounts of the full-length 3x-Flag-tagged CNF_Y_ protein (25, 50, 100, 250, and 500 µg/ml bacterial extract of bacteria overexpressing the toxin derivatives) at 4°C for 1 h. Subsequently, whole cell extracts were prepared and bound CNF_Y_ proteins were detected by Western blotting using a monoclonal anti-FLAG antibody.

